# Mitochondria-targeted hydrogen sulfide donor reduces atherogenesis by reprogramming macrophages and increasing UCP1 expression in vascular smooth muscle cells

**DOI:** 10.1101/2024.05.15.594319

**Authors:** Aneta Stachowicz, Anna Wiśniewska, Klaudia Czepiel, Bartosz Pomierny, Alicja Skórkowska, Beata Kuśnierz-Cabala, Marcin Surmiak, Katarzyna Kuś, Mark E. Wood, Roberta Torregrossa, Matthew Whiteman, Rafał Olszanecki

## Abstract

**Aims:** Atherosclerosis is a leading cause of morbidity and mortality in the Western countries. A growing body of evidence points to the role of mitochondrial dysfunction in the pathogenesis of atherosclerosis. Recently, it has been shown that mitochondrial hydrogen sulfide (H_2_S) can complement the bioenergetic role of Krebs cycle leading to improved mitochondrial function. However, controlled, direct delivery of H_2_S to mitochondria was not investigated as a therapeutic strategy in atherosclerosis. Therefore, the aim of our study was to comprehensively evaluate the influence of prolonged treatment with mitochondrial H_2_S donor AP39 on the development of atherosclerotic lesions in apolipoprotein E knockout (apoE^-/-^) mice.

**Results:** Our results indicated that AP39 reduced atherosclerosis in apoE^-/-^ mice and stabilized atherosclerotic lesions through decreased total macrophage content and increased collagen depositions. Moreover, AP39 reprogrammed macrophages from proinflammatory M1 to anti-inflammatory M2 in atherosclerotic lesions. It also upregulated pathways related to mitochondrial function, such as cellular respiration, fatty acid β-oxidation and thermogenesis while downregulated pathways associated with immune system, platelet aggregation and complement and coagulation cascades in the aorta. Furthermore, treatment with AP39 increased the expression of mitochondrial brown fat uncoupling protein 1 (UCP1) in vascular smooth muscle cells (VSMCs) in atherosclerotic lesions and upregulated mRNA expression of other thermogenesis-related genes in the aorta but not perivascular adipose tissue (PVAT) of apoE^-/-^ mice. Finally, AP39 treatment decreased markers of activated endothelium and increased endothelial nitric oxide synthase (eNOS) expression and activation.

**Conclusions:** Taken together, mitochondrial H_2_S donor AP39 could provide potentially a novel therapeutic approach to the treatment/prevention of atherosclerosis.

## Introduction

Atherosclerosis is a leading cause of morbidity and mortality in the Western countries. It is a complex inflammatory disease of arteries that is characterized by lipid-rich necrotic core and rupture-prone fibrous cap. The rupture of the atherosclerotic plaque can cause life-threatening complications, including coronary artery disease, stroke, peripheral artery disease and myocardial infarction ^1^. Despite the high prevalence of the disease, numerous aspects of pathogenesis remain unclear. It has been recently proposed that mitochondrial dysfunction contributes to the initiation and progression of atherosclerosis. Dysfunctional mitochondria produce excessive reactive oxygen species (ROS) through the electron transport chain (ETC), activate apoptosis, and initiate the development of inflammation by NLRP3 inflammasome activation ^2^. Importantly, mitochondrial dysfunction, which leads to reduced energy supply in cells, affects endothelial function, vascular smooth muscle cell (VSMC) proliferation and apoptosis, and macrophage polarization ^3^. Several studies have shown that mitochondrial metabolism can influence macrophage reprogramming ^4^. Proinflammatory M1 macrophages use glycolysis as the main energy source, thus inhibiting mitochondrial function. In contrast, anti-inflammatory M2 macrophages rely on oxidative phosphorylation as the major energy source, hence mitochondrial function is required for the maintenance of M2 phenotype ^5^. Thus, improving the mitochondrial function may contribute to atherosclerosis regression by reprogramming mitochondrial metabolism of macrophages towards anti-inflammatory phenotype.

Growing body of evidence suggest that hydrogen sulfide (H_2_S) plays a protective role in the pathogenesis of atherosclerosis by inhibiting inflammation, monocyte adhesion to endothelium, macrophage foam cell formation, VSMC proliferation and platelet aggregation ^6^. H_2_S is a small gaseous signaling molecule that regulates vascular tone, inflammation, proliferation and cell death ^7^. Endogenous H_2_S is produced by cystathionine-β-synthase (CBS), cystathionine-γ-lyase (CSE) and 3-mercaptopyruvate sulfurtransferase (3-MST). CBS and CSE are cytosolic enzymes, whereas 3-MST is both a mitochondrial and cytosolic enzyme. It is well-known that higher concentrations of H_2_S exhibit inhibitory impact on complex IV of ETC ^8^. However, it has been recently demonstrated that H_2_S at low concentrations has a physiological function in supporting cellular bioenergetics. H_2_S derived from 3-MST was shown to serve as an inorganic source of energy and electron donor to complex II of ETC. Thus, intramitochondrial H_2_S could complement the bioenergetic role of Krebs cycle as an alternative source of electrons for oxidative phosphorylation and ATP production leading to improved mitochondrial function ^9^. However, controlled, direct delivery of H_2_S to mitochondria was not considered as a therapeutic strategy in the treatment of atherosclerosis.

Recently, a mitochondria-targeted, slow-releasing H_2_S donor named AP39 has been synthesized. AP39 consists of H_2_S donating moiety linked to triphenylphosphonium (TPP^+^) – a mitochondria targeting motif. It has been demonstrated that AP39 at lower concentrations stimulated cellular bioenergetics and reduced oxidative stress ^10^. Of note, the ability of AP39 to stimulate cellular bioenergetics and improve mitochondrial function could play a protective role in the development of atherosclerosis. Thus, the aim of our study was to comprehensively investigate the influence of prolonged treatment with mitochondrial donor of H_2_S AP39 on the development of atherosclerotic lesions in apolipoprotein E knockout (apoE^-/-^) mice on a high-fat diet (HFD). ApoE^-/-^ mice spontaneously develop atherosclerotic lesions, hypercholesterolemia, and dyslipidemia, and hence are a popular animal model of atherosclerosis.

## Materials and Methods

### Animal studies

Twenty-six female apoE^−/−^ mice on the C57BL/6J background (B6.129P2-ApoEtm1Unc/J) were obtained from the Jackson Laboratory (USA). The animals were maintained on 12 h dark/12 h light cycles at room temperature (22.5 ± 0.5 °C) and 45-55% humidity with access to water ad libitum and diet. At the age of 8 weeks the mice were fed with an HFD (Sniff, Germany: E15122-34 containing 10% fat and 95 mg/kg cholesterol) for 16 weeks. The animals were divided into two groups: female apoE^−/−^ mice on an HFD (control) (*n* = 13) treated with vehicle (10% DMSO) and female apoE^−/−^ mice on an HFD treated with AP39 (*n* = 13). Sample size was calculated in Statistica software based on our previous results ^11^. AP39 was synthesized in-house as previously described ^12^ and administered subcutaneously to the mice at a dose of 0.1 mg/kg of body weight per day three days a week for 16 weeks. The dose of AP39 was selected based on previous experiments ^13^. At the age of 6 months the mice were euthanized 5 min after injection of Fraxiparine i.p (1000 UI; Sanofi-Synthelabo France) in chamber filled with carbon dioxide at a rate of 20-30% CO_2_ chamber volume per minute, in accordance with AVMA Panel 2007 recommendations and institutional IACUC guidelines. The selected tissues (aortas, hearts, perivascular adipose tissue (PVAT)) were dissected, and the blood was collected. 7 aortas (PVATs) in each group were collected for proteomics experiments and 6 aortas (PVATs) in each group for RT‒qPCR experiments. All animal procedures were conformed with the guidelines from Directive 2010/63/EU of the European Parliament on the protection of animals used for scientific purposes and were approved by the Jagiellonian University Ethical Committee on Animal Experiments (No. 517/2021).

### Atherosclerotic lesion assessment

The development of atherosclerotic lesions was evaluated using cross section method. Hearts with ascending aorta were embedded in OCT compound (CellPath, UK), snap frozen and sectioned (10 μm thickness) for histological and immunohistochemical analysis. The aortic sections were stained with Oil Red-O (Sigma-Aldrich, USA) to measure the area of atherosclerotic plaques. Furthermore, hematoxylin-eosin (H&E) and Masson’s trichrome staining were performed to assess the presence of necrotic cores and collagen deposition within the lesions. The necrotic core area was defined as area without H&E staining within the lesion and was shown as a percentage of the total plaque area. Masson’s trichrome staining was expressed as a percentage of collagen area (blue area) relative to the total plaque area. Aortic images were captured using Olympus BX50 (Olympus, Japan) microscope and the data were analyzed by the LSM Image Browser software (Zeiss, Germany).

### Immunohistochemical staining of aortic roots

Frozen sections of ascending aorta were stained with primary antibodies against CD68 (Serotec, UK) and smooth muscle α-actin (SMA) (Sigma-Aldrich, USA) (dilution 1:800). To access macrophage polarization, antibodies against F4/80 (dilution 1:100, Abcam UK), nitric oxide synthase 2 (iNOS) (dilution 1:200, Abcam UK) and arginase 1 (dilution 1:50, Cell Signaling, USA) were used. To evaluate uncoupling protein 1 (UCP1) expression in VSMCs, antibodies against the marker of VSMCs - calponin (dilution 1:200, Proteintech, USA) and UCP1 (dilution 1:200, Thermo Scientific USA) were used. All sections were mounted in anti-fade mounting medium with DAPI (Vector Laboratories, USA) to reveal nuclear staining. Sections were scanned using Leica Stellaris 8 WLL, DLS confocal microscope (Leica, Germany) and analyzed using LAS X software (Leica, Germany) with the co-expression module. Each section was scanned with 40x, NA 0.8 objective, at three locations (macrophage polarization) or with 20x, NA 0.75 objective (UCP1 expression). Cells with detected co-localization of three fluorescent signals from DAPI (nuclei), F4/80 (total macrophages), and iNOS/arginase 1 (M1/M2 phenotype) were counted and divided by the number of total macrophages. To evaluate UCP1 expression in VSMCs, cells with detected co-localization of three fluorescent signals from DAPI (nuclei), calponin (VSMCs), and UCP1 were counted and divided by the area of atherosclerotic lesions.

### Biochemical measurements

The blood was centrifuged at 1000 × g at 4°C for 10 min and the plasma was collected and stored at - 80°C. The levels of total cholesterol, triglycerides (TG), low-density lipoproteins (LDL) and high-density lipoproteins (HDL) were measured using an enzymatic method on a Cobas 8000 analyzer (Roche Diagnostics, USA). Plasma concentrations of some markers related to inflammation, coagulation and angiogenesis were determined using the custom-made xMAP technology Luminex assays (R&D Systems, USA) and the Luminex MAGPIX System (Luminex Corp., USA). The ratio of reduced glutathione (GSH) to oxidized glutathione (GSSG) was measured by commercially available Glutathione Colorimetric Detection Kit (Invitrogen, USA).

### Flow cytometry

Peritoneal cells were harvested by injecting 5 mL of PBS into the peritoneal cavity. The cells were centrifuged, washed, and counted. Equal numbers of cells were first stained with BD Horizon Fixable Viability Stain 450 (BD Biosciences, USA) to exclude dead cells. Subsequently, these cells were preincubated with anti-mouse CD16/CD32 antibody to block FcγRII/III receptors and were labelled with PerCP-conjugated anti-mouse F4/80, FITC-conjugated anti-mouse/human CD11b and APC-conjugated anti-mouse CD206 antibodies (BioLegend, USA). After surface staining, the cells were fixed and permeabilized using BD Cytofix/Cytoperm buffer (BD Biosciences, USA) and then stained with PE-Cyanine7-coniugated anti-mouse iNOS (eBioscience, USA). Unstained cells were used to establish flow cytometer settings. The samples were acquired on a FACSCanto II flow cytometer (BD Biosciences, USA) and analyzed using FACSDiva software. The gating strategy used excluded doublets and dead cells from the analysis, and additional gates were set based on fluorescence minus one (FMO) control. F4/80 and CD11b were used as pan-macrophage markers, while iNOS and CD206 were used as markers of M1 and M2 macrophages, respectively. F4/80-positive/CD11b-positive/iNOS-positive/CD206-negative cells were defined as M1 macrophages, while F4/80-positive/CD11b-positive/iNOS-negative/CD206-positive cells were defined as M2 macrophages.

### Liquid chromatography-tandem MS (LC‒MS/MS) analysis of mouse aorta

Mouse aorta was homogenized using a Tissue Lyser LT (Qiagen, Germany) and lysed in a buffer containing 0.1 M Tris-HCl, pH 7.6, 2% SDS, and 50 mM DTT (Sigma Aldrich, USA) at 96 °C for 10 min. Seventy micrograms of protein were digested overnight using the filter-aided sample preparation (FASP) method with Trypsin/Lys-C mix (Promega, USA). Next, the samples were purified with C18 Ultra-Micro Spin Columns (Harvard Apparatus, USA). For mouse aorta spectral library preparation, equal amounts of peptides from all samples were subjected to a high-pH fractionation protocol on C18 Micro Spin Columns (Harvard Apparatus, MA).

One microgram of peptide was injected into a nanoEase^TM^ M/Z Peptide BEH C18 75 µm i.d. × 25 cm column (Waters, USA) via a nanoEase^TM^ M/Z Symmetry C18 180 µm i.d. × 2 cm trap column (Waters, USA) and separated with a flow rate of 250 nL/min on an UltiMate 3000 HPLC system coupled to a Orbitrap Exploris™ 480 Mass Spectrometer (Thermo Scientific, USA). For data-dependent (DDA) acquisition, spectra were collected for 145 min in full scan mode (350–1400 Da) with fixed cycle time (1.3s) and AGC set to 200%. For data-independent (DIA) acquisition, spectra were collected for 145 min in full scan mode (400–1250 Da), followed by 55 DIA scans using a variable precursor isolation window approach and AGC set to custom 1000%.

DDA MS data were searched against the mouse UniProt database and MaxQuant Contaminants list using the Pulsar search engine in Spectronaut software (Biognosys, Switzerland) with the following parameters: ± 40 ppm mass tolerance on MS1 and MS2 levels, mutated decoy generation method, Trypsin/Lys-C enzyme specificity, 1% protein and PSM false discovery rate (FDR). The generated mouse aorta library was used to analyse DIA MS data in Spectronaut software. MS data were filtered by 1% FDR at the peptide and protein levels, while quantitation was performed at the MS2 level without imputation. Statistical analysis of differential protein abundance was performed at both the MS1 and MS2 levels using unpaired t tests with multiple testing correction after Storey. Functional grouping and pathway analysis were performed using PINE (Protein Interaction Network Extractor) software ^14^ with the STRING and GeneMANIA databases using a score confidence of 0.4 and a ClueGO p value cutoff < 0.05. The mass spectrometry data have been deposited to the ProteomeXchange Consortium via the PRIDE partner repository ^15^ with the dataset identifier PXD050076.

### Western blot analysis

Mouse aorta and PVAT were lysed as described in proteomics section. Protein samples were separated on SDS-polyacrylamide gels (7.5% or 12%) using a Laemmli buffer system and then semidry transferred to nitrocellulose membranes by a Trans-Blot Turbo Transfer System (Bio-Rad, USA). The membranes were blocked with 5% bovine serum albumin in PBS at room temperature for 1 h and incubated overnight at 4 °C with specific anti-phospho-eNOS (Ser1177), anti-eNOS (Cell Signaling, USA), anti-ACAA2 (Sigma‒Aldrich, USA), anti-UCP1 (Invitrogen, USA) and anti-COX IV (MyBioSource, USA) (concentration 1:1000) primary antibodies. Incubation with HRP-conjugated secondary antibodies (GE Healthcare, USA) was performed at room temperature for 1 h (dilution 1:10 000). Protein bands were developed by Clarity™ Western ECL Substrate (Bio-Rad, USA) for 5 min. Protein bands were visualized and imaged by an ImageQuant LAS 500 scanner (GE Healthcare, USA). After transfer, the blots were stained with Ponceau S solution (Sigma‒Aldrich, USA) and developed for total protein level normalization. Data were analyzed in the Image Studio^TM^ Lite software (LI-COR Biosciences, USA).

### Quantitative reverse transcription polymerase chain reaction (RT‒qPCR)

Briefly, total RNA was isolated from the aortas using the RNeasy Fibrous Tissue Mini Kit (Qiagen, Germany), according to the manufacturer’s instructions. cDNA was synthesized by the reverse transcription of total RNA, using a High-Capacity Reverse Transcription Kit (Applied Biosystems, USA). The primers used in these experiments included primers purchased from Bio-Rad (*Nos3*, *Nrf2*, *Pparg*, *Gapdh*), RealTimePrimers (*Pgc1, Rpl13a*) as well as following primers *Cidea*: 5’ TGCTCTTCTGTATCGCCCAGT 3’ (forward), 5’ GCCGTGTTAAGGAATCTGCTG 3’ (reverse); *Elovl3*: 5’ TCCGCGTTCTCATGTAGGTCT 3’ (forward), 5’ GGACCTGATGCAACCCTATGA 3’ (reverse). 2x SsoAdvanced™ Universal SYBR® Green Supermix (Bio-Rad, USA) was used to carry out real-time PCR reaction. Analysis of relative gene expression was performed by the CFX96 Touch Real-Time PCR Detection System (Bio-Rad,USA) with *Gapdh* and *Rpl13a* as internal reference genes for aorta and PVAT, respectively. Data were analyzed using the 2–ΔΔCt method in an Excel spreadsheet.

### Statistical analysis

Data are expressed as the mean ± SEM. The equality of variance and normality of the data were checked by the F test and Shapiro‒Wilk test, respectively. Unpaired t test or unpaired t test with Welch’s correction were used to analyze differences between control and AP39 group (GraphPad Prism 9.3.1, USA). Values of p < 0.05 were considered statistically significant.

## Results

### AP39 reduced atherogenesis in apoE^-/-^ mice

In our experiments, we used AP39, a chemical compound that increases the levels of H_2_S within mitochondria. AP39 was administered subcutaneously to apoE^-/-^ mice on HFD three days a week in a dose of 0.1 mg/kg of body weight per day for 16 weeks. The mean weight of mice did not differ between control mice and AP39-treated mice over 16 weeks of experiment duration as well as at the end of experiment. Moreover, the food consumption rate was similar between control mice and AP39-treated mice over 16 weeks of experiment (Supplemental Figure 1).

To evaluate the impact of AP39 on the development of atherosclerosis, we measured the size of atherosclerotic lesions in the aorta of apoE^-/-^ mice by cross section method. AP39 treatment significantly decreased atherosclerotic plaque in the aorta of apoE^-/-^ mice (Fig. 1A). The reduction of atherogenesis by AP39 was not associated with the alterations in total cholesterol, HDL, LDL and TG in the plasma of apoE^-/-^ mice (Table 1).

**Table 1.**
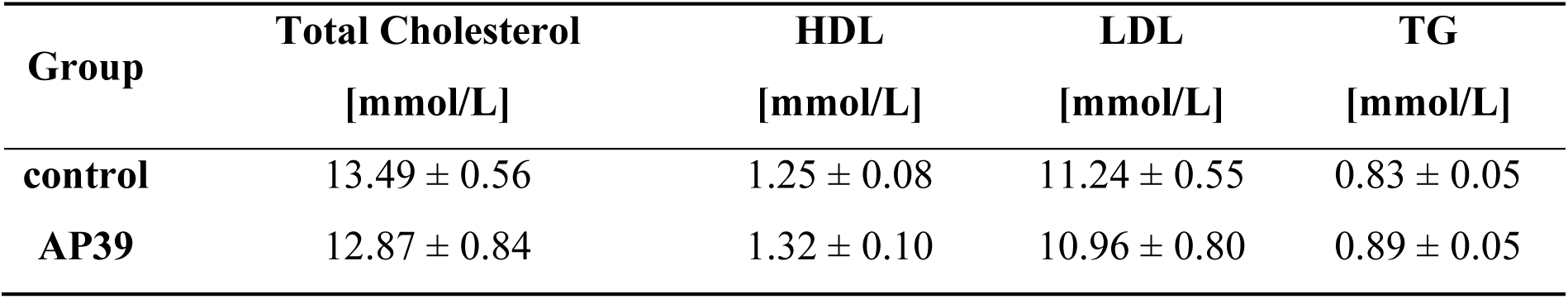
Plasma levels of total cholesterol, HDL, LDL, and TG in control and AP39-treated mice, presented as mean ± SEM; n = 7 per group.

| Group | Total Cholesterol<br>[mmol/L] | HDL<br>[mmol/L] | LDL<br>[mmol/L] | TG<br>[mmol/L] |
| --- | --- | --- | --- | --- |
| control | 13.49 ± 0.56 | 1.25 ± 0.08 | 11.24 ± 0.55 | 0.83 ± 0.05 |
| AP39 | 12.87 ± 0.84 | 1.32 ± 0.10 | 10.96 ± 0.80 | 0.89 ± 0.05 |

In addition, AP39 administration did not significantly impact the plasma levels of markers related to inflammation (ICAM-1, M-CSF, MMP-9), coagulation (PAI-1, P-selectin) and angiogenesis (VEGF, ANG-2) (Table 2).

**Table 2.** Plasma levels of markers related to inflammation, coagulation, and angiogenesis in control and AP39-treated apoE^-/-^ mice, presented as mean ± SEM; n = 7 per group.

| Plasma marker [pg/mL] | control | AP39 |
| --- | --- | --- |
| ICAM-1 | 27241 ± 5727 | 30532 ± 4777 |
| M-CSF | 7.96 ± 2.10 | 8.87 ± 1.62 |
| MMP-9 | 3325 ± 1020 | 3203 ± 591.7 |
| PAI-1 | 652.4 ± 85.2 | 738.3 ± 96.19 |
| P-selectin | 22241 ± 2947 | 20581 ± 3671 |
| VEGF | 71.39 ± 5.07 | 74.55 ± 5.56 |
| ANG-2 | 8247 ± 650.4 | 10361 ± 780.3 |

**Figure 1.**
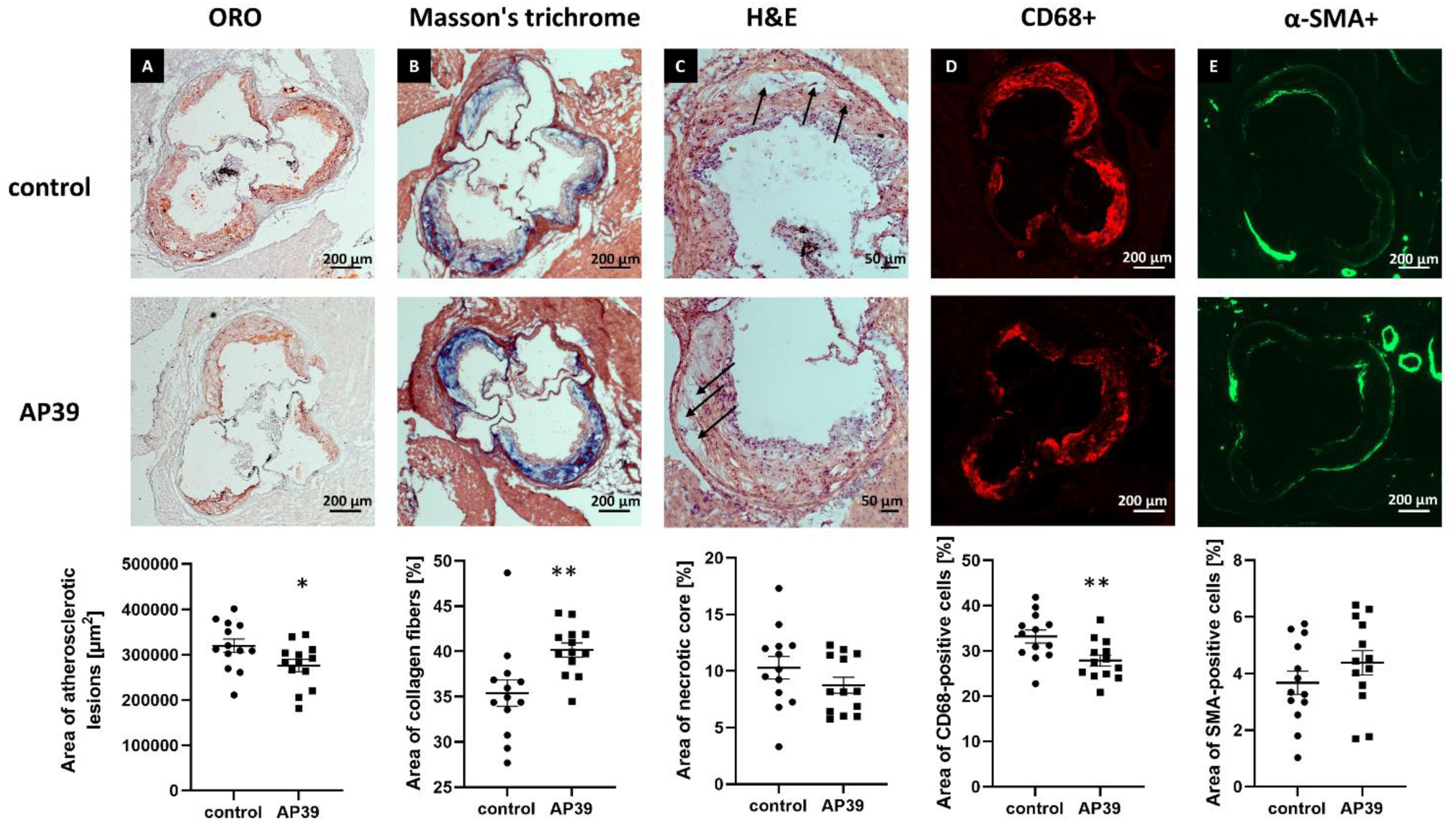
AP39 reduces atherogenesis and stabilizes atherosclerotic plaque in apoE^-/-^ mice. Representative micrographs showing oil-red O – stained atherosclerotic lesions **(A)**, Masson’s trichrome stained collagen depositions **(B)** and HE stained necrotic cores **(C)** in the aorta of control and AP39-treated mice. Necrotic cores are indicated by black arrows. Immunohistochemical staining of aortic roots showing CD68-positive macrophages **(D)** and SMA **(E)** in control and AP39-treated mice. Mean ± SEM; *p<0.05, **p<0.01 compared to control mice; n=13; unpaired two-tailed Student’s *t*-test.

### AP39 stabilized atherosclerotic plaque by decreasing proinflammatory M1 macrophages and increasing anti-inflammatory M2 macrophages

To evaluate the impact of mitochondrial donor of H_2_S AP39 on plaque stability, we measured the area of collagen deposition, necrotic core as well as macrophage and SMA content within lesions. Our data showed that AP39 administration caused the stabilization of atherosclerotic lesions in apoE^-/-^ mice. It significantly increased collagen depositions (Fig. 1B) as measured by Masson’s trichrome staining and decreased macrophage content as evidenced by CD68 staining (Fig. 1D) in atherosclerotic plaque of apoE^-/-^ mice. AP39 treatment did not change the area of necrotic core (Fig. 1C) as well as SMA content (Fig. 1E).

**Figure 2.**
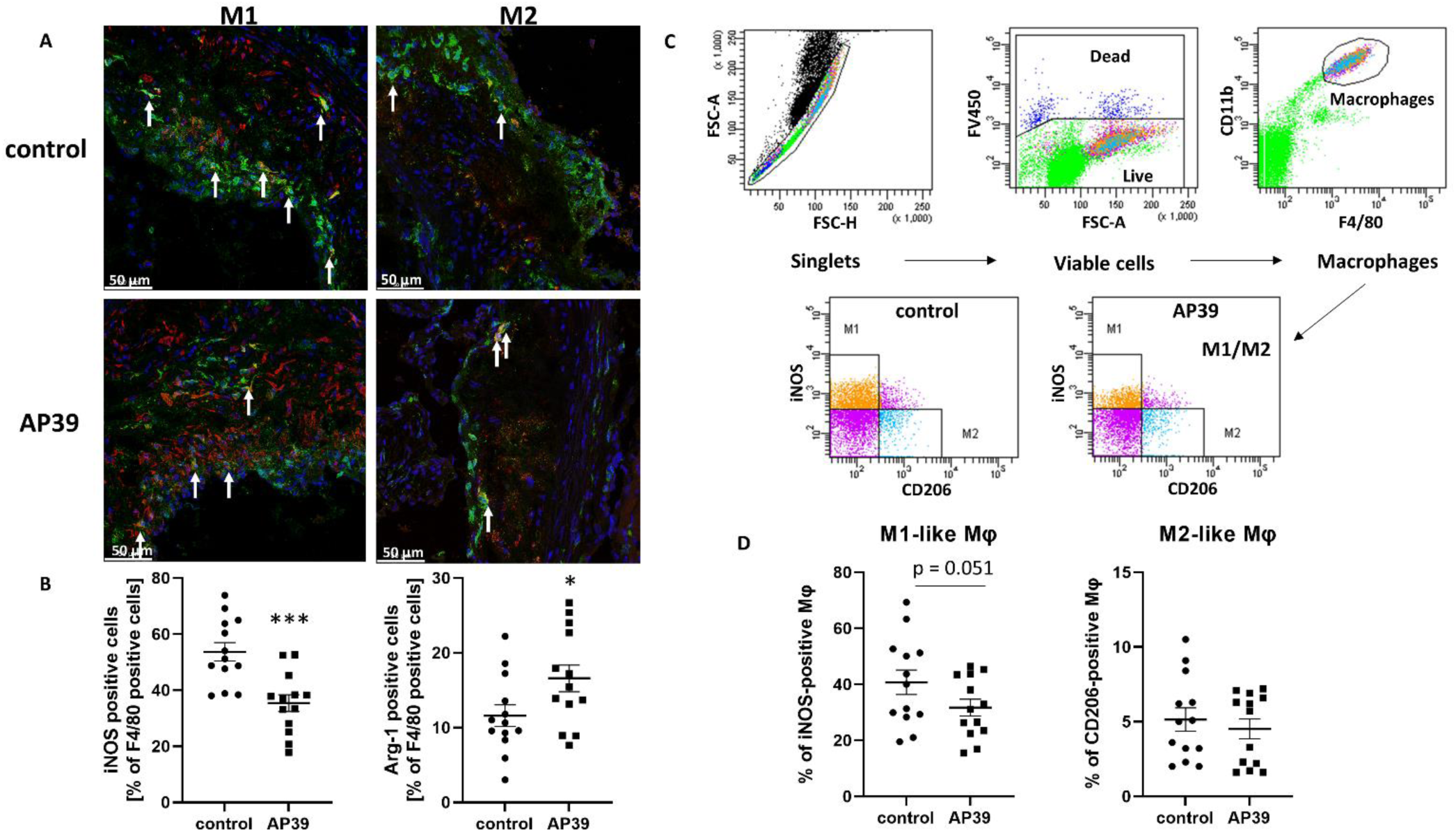
AP39 decreases polarization of macrophages to proinflammatory M1 phenotype and increases polarization to anti-inflammatory M2 phenotype. Representative immunohistochemical staining of aortic roots showing F4/80 (green), DAPI (blue) and iNOS (red) (M1 phenotype) or arginase 1 (red) (M2 phenotype) co-localization **(A)** in control and AP39-treated mice. White arrows indicate M1 or M2 macrophages. Quantitative analysis of M1 and M2 macrophage phenotypes in atherosclerotic lesions **(B)**. Flow cytometry phenotyping of peritoneal macrophages in vivo. Gating strategy indicating the procedure to quantify M1-like (CD11b/F4/80/iNOS-positive) and M2-like (CD11b/F4/80/CD206-positive) macrophages in the peritoneum of control and AP39-treated mice **(C)**. Quantitative analysis of M1 and M2 macrophage phenotypes in peritoneum **(D)**. Mean ± SEM; *p<0.05 ***p<0.001 compared to control mice; n=13; unpaired two-tailed Student’s *t*-test.

To further explore the impact of AP39 on plaque stabilization, we investigated whether AP39 could change the content of proinflammatory M1 and anti-inflammatory M2 macrophages within atherosclerotic lesions. Interestingly, treatment with AP39 decreased the level of M1 macrophages (Fig. 2A and B, Supplemental Figure 2A) in atherosclerotic lesions of apoE^-/-^ mice as well as increased the content of M2 macrophages (Fig. 2A and B, Supplemental Figure 2B). We also measured the level of M1 and M2 macrophages among peritoneal macrophages isolated from AP39-treated mice and control mice using flow cytometry. Treatment with AP39 tended to reduce the number of proinflammatory M1 macrophages but not M2 macrophages (Fig. 2C and D) in the peritoneum of apoE^-/-^ mice.

### Proteomic analysis revealed pathways downregulated by AP39 in the aorta, including pathways related to immune system, platelet activation and aggregation as well as complement and coagulation cascades

To comprehensively investigate the changes in protein expression upon AP39 administration, a quantitative proteomics analysis was conducted. In total, 6565 protein groups across all biological conditions were identified and quantified. A summary of the quality control for the LC‒MS/MS runs is shown in Supplemental Figure 3. We found a total of 1048 proteins altered in the aorta of AP39-treated mice compared to control group (Fig. 3A). The detailed list of differentially expressed proteins and their fold changes across all biological conditions are presented in Supplemental Table 1.

To identify the functional pathways and biological networks, we performed enriched pathway analysis based on protein expression using PINE software. Pathways related to immune system, platelet activation and aggregation, and complement and coagulation cascades were downregulated upon AP39 treatment, shown as blue central nodes (Fig. 3B). Among others, AP39 administration decreased expression of thromboxane-A synthase (TXA), a potent inducer of blood vessel constriction and platelet aggregation, in the aorta of apoE^-/-^ mice (Supplemental Table 1). On the other hand, pathways associated with mitochondrial function, such as Krebs cycle, oxidative phosphorylation, thermogenesis, and mitochondrial fatty acid β-oxidation, shown as orange central nodes, were upregulated because of AP39 administration (Fig. 4A).

Interestingly, some of our proteomic results in the aorta were in line with the observed stabilization of atherosclerotic plaque upon AP39 administration in apoE^-/-^ mice. We showed increased expression of different collagen types, such as collagen alpha-1(I) chain (COL1A1) as well as decreased expression of different matrix metalloproteinases (MMPs), including 72 kDa type IV collagenase (MMP-2), macrophage metalloelastase (MMP-12), matrix metalloproteinase-19 (MMP-19), and stromelysin-1 (MMP-3) in the aorta of AP39-treated mice (Supplemental Table 1). Of note, consistently with immunohistochemistry data in atherosclerotic lesions, our proteomic approach also pointed out to the decreased expression of macrophage and monocyte markers in the aorta of AP39-treated mice. Among others, expression of CD68 antigen, macrophage-expressed gene 1 protein (MPEG1) and monocyte differentiation antigen CD14 was downregulated upon AP39 administration in the aorta of apoE^-/-^ mice (Supplemental Table 1).

Importantly, treatment with AP39 resulted in the beneficial changes of several proteins involved in the pathogenesis of atherosclerosis. Expression of proteins participated in the activation of endothelium, such as P-selectin and vascular cell adhesion protein 1 (VCAM-1) was downregulated in the aorta of AP39-treated mice. In addition, upon AP39 administration we observed decreased expression of different subunits of neutrophilic type NADPH oxidase, a main source of reactive oxygen species (ROS) during atherogenesis. Moreover, expressions of proteins regulating macrophage polarization, such as dipeptidyl peptidase 1 (DPPI), dipeptidyl peptidase 9 (DPP9), glia maturation factor gamma (GMFG) and lipocalin-2 (LCN2) were downregulated in the aorta of AP39-treated mice (Supplemental Table 1).

**Figure 3.**
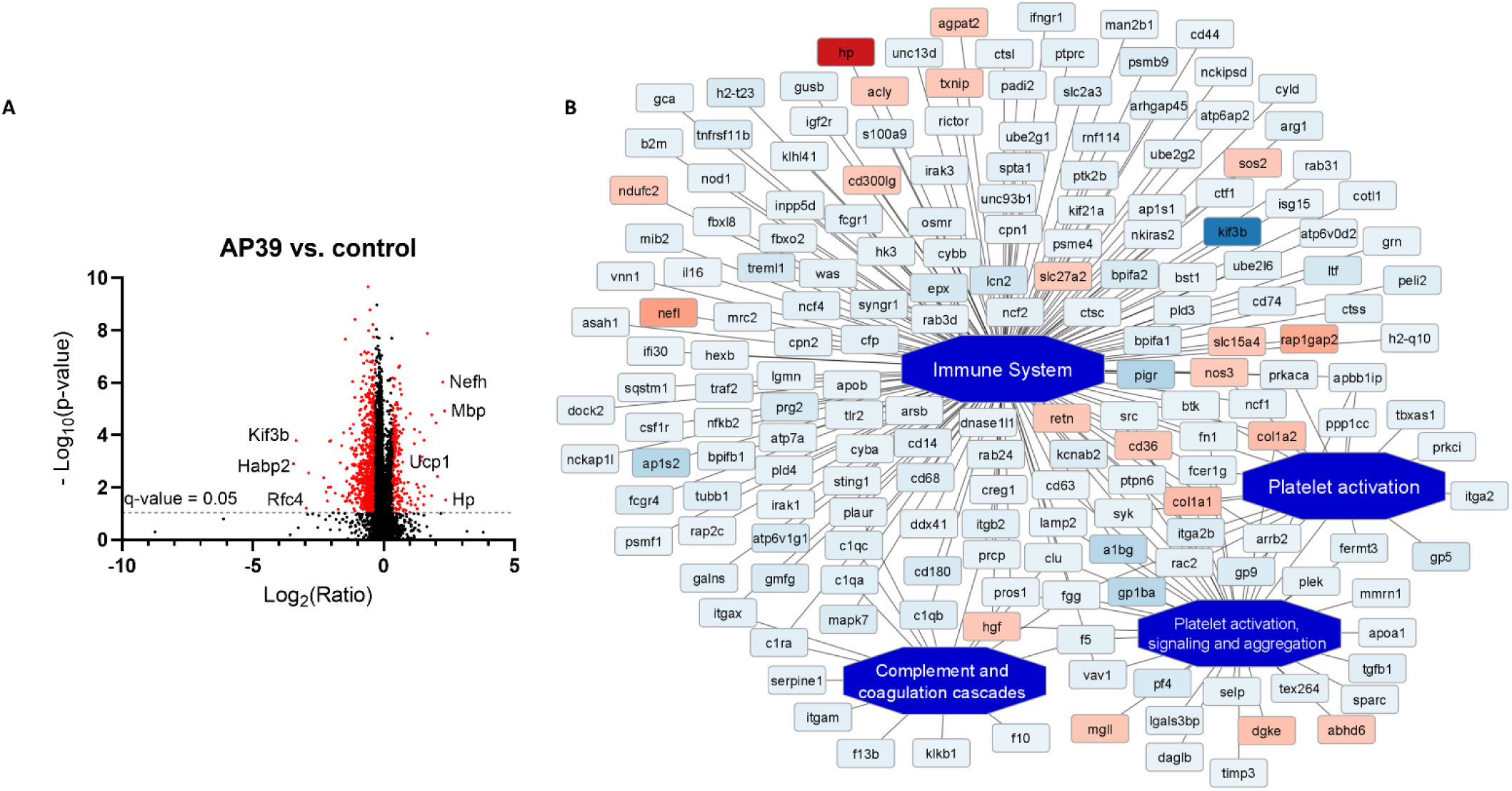
Proteomic analysis of the aorta reveals downregulated pathways related to immune system, platelet activation and aggregation as well as complement and coagulation cascades upon treatment with AP39. Volcano plot of differentially expressed proteins showing the log2 ratio of protein expression versus -log10 p value in AP39 group compared to control **(A)**. The most upregulated and downregulated proteins are shown. Enriched functional network in AP39-treated mice compared to control, as generated by PINE **(B)**. Activated pathways are shown as orange central nodes, and inhibited pathways are shown as blue central nodes along with red (upregulated) or blue (downregulated) protein nodes. q-value<0.05; n=7; unpaired two-tailed Student’s *t*-test.

### AP39 upregulated pathways related to mitochondrial function, such as thermogenesis, cellular respiration and fatty acid β-oxidation

Our proteomic analysis indicated that treatment with AP39 led to the significant upregulation of proteins engaged in cellular respiration: oxidative phosphorylation and Krebs cycle. AP39 administration increased expression of all electron transport chain proteins, including Complex I (different subunits of NADH dehydrogenase), Complex II (succinate dehydrogenase flavoprotein subunit (SDHA) and iron-sulfur subunit (SDHB), Complex III (different subunits of cytochrome b-c1 complex) and Complex IV (different subunits of cytochrome c oxidase, e.g., subunit 4 isoform 1 (COX IV)). Furthermore, proteins involved in Krebs cycle (e.g., citrate synthase (CS), isocitrate dehydrogenase subunit alpha (IDH3A) and gamma 1 (IDH3G), aconitate hydratase (ACO2), pyruvate dehydrogenase E1 component subunit beta (PDHB)) were upregulated upon AP39 treatment in the aorta of apoE^-/-^ mice (Supplemental Table 1).

Interestingly, AP39 administration increased expression of proteins participated in thermogenesis, such as mitochondrial brown fat uncoupling protein 1 (UCP1), carnitine O-palmitoyltransferase 2 (CPT2), 1-acylglycerol-3-phosphate O-acyltransferase ABHD5 (ABHD5), and hormone-sensitive lipase (HSL) in the aorta of apoE^-/-^ mice (Supplemental Table 1). Moreover, AP39 administration also upregulated expression of proteins responsible for mitochondrial fatty acid β-oxidation, including 3-ketoacyl-CoA thiolase (ACAA2), short-chain specific acyl-CoA dehydrogenase (SCAD), and trifunctional enzyme subunit alpha (HADHA) and beta (HADHB) (Supplemental Table 1).

To confirm increased expression of proteins responsible for cellular respiration and mitochondrial fatty acid β-oxidation, we performed Western blot analysis of selected proteins: COX IV and ACAA2. Indeed, AP39 administration led to the upregulation of COX IV (Fig. 4B and D) and ACAA2 (Fig. 4B and C) in the aorta of apoE^-/-^ mice. Uncropped Western blot images of COX IV and ACAA2 are presented in Supplemental Figure 4.

**Figure 4.**
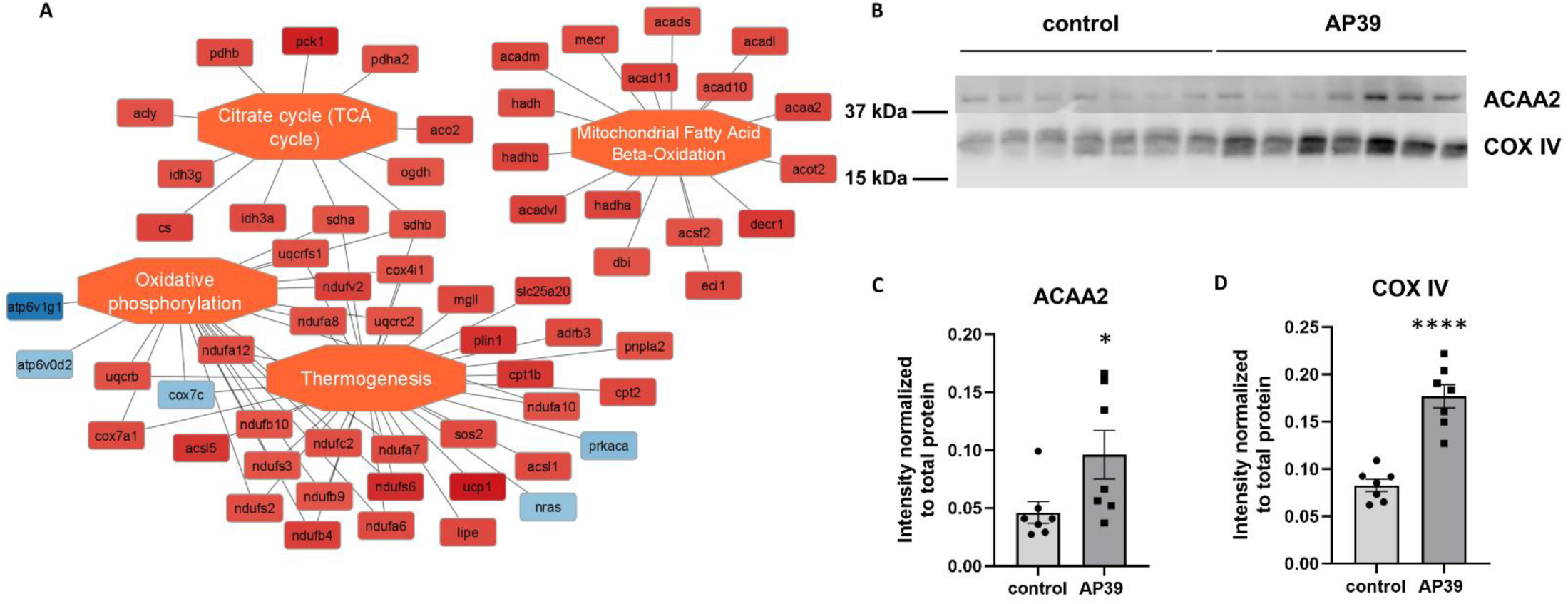
AP39 increases thermogenesis, cellular respiration, and β-oxidation in the aorta of apoE^-/-^ mice. Enriched functional network of mitochondrial processes in AP39-treated mice compared to control, as generated by PINE **(A)**. Activated pathways are shown as orange central nodes, and inhibited pathways are shown as blue central nodes along with red (upregulated) or blue (downregulated) protein nodes. Western blot of ACAA2 and COX IV in the aorta of control and AP39-treated mice **(B)**. Quantitative analysis of Western blots for ACAA2 (**C**) and COX IV **(D)** expression in the aorta of control and AP39-treated mice. Mean ± SEM; *p<0.05, ****p<0.0001; n=7; unpaired two-tailed Student’s *t*-test.

### AP39 increased UCP1 expression and thermogenesis-related gene expression in the aorta but not PVAT

Our proteomic data pointed to the upregulated expression of UCP1, and other proteins involved in thermogenesis upon AP39 administration in the aorta of apoE^-/-^ mice. To confirm increased expression of UCP1, a main regulator of thermogenesis, we performed Western blot analysis in the aorta and PVAT of apoE^-/-^ mice. Indeed, AP39 administration led to the upregulation of UCP1 (Fig. 5A and B) in the aorta but not PVAT (Fig. 5A and C). Uncropped Western blot images of UCP1 are presented in Supplemental Figure 4 and 5.

To further explore the impact of AP39 on thermogenesis pathway, we investigated the expression of thermogenesis-related genes in the aorta and PVAT of apoE^-/-^ mice. Interestingly, we observed increased mRNA expression of cell death inducing DFFA like effector A (*Cidea*), peroxisome proliferator activated receptor gamma (*Pparg*), peroxisome proliferator-activated receptor gamma coactivator 1-alpha (*Pgc1*), and nuclear factor erythroid 2-related factor 2 (*Nrf2*) in the aorta (Fig. 5D) but not PVAT (Fig. 5E) of apoE^-/-^ mice. mRNA expression of ELOVL fatty acid elongase 3 (*Elovl3*) also tended to increase in the aorta (Fig. 5D) but not PVAT (Fig. 5E).

**Figure 5.**
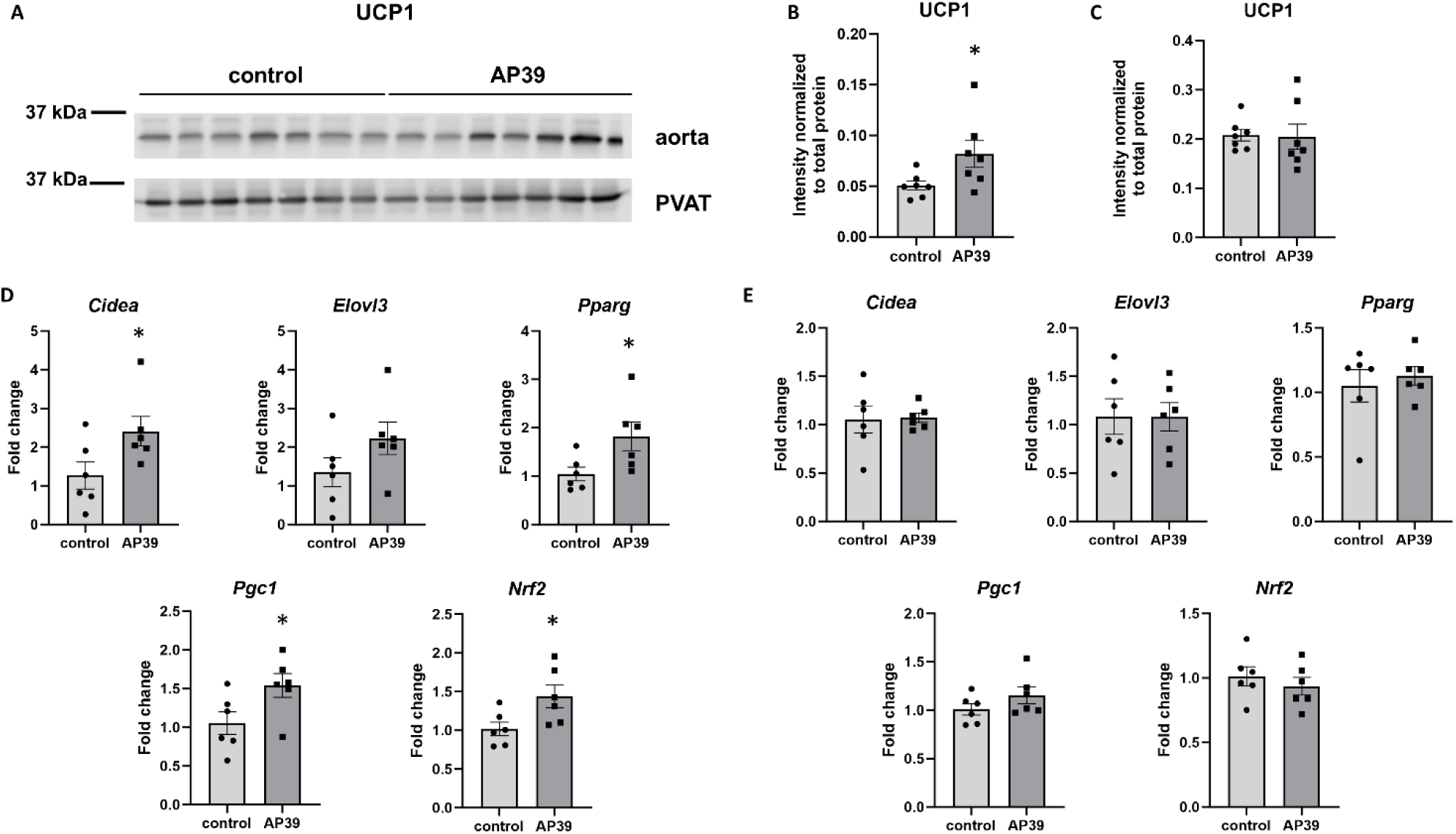
AP39 increases UCP1 expression and thermogenesis-related gene expression in the aorta but not PVAT of apoE^-/-^ mice. Western blot of UCP1 in the aorta and PVAT of control and AP39-treated mice **(A)**. Quantitative analysis of Western blots for UCP1 expression in the aorta **(B)** and PVAT **(C)** of control and AP39-treated mice. mRNA expression of thermogenesis-related genes including *Cidea*, *Elovl3*, *Pparg*, *Pgc1* and *Nrf2* in the aorta (**D**) and PVAT (**E**) of control and AP39-treated mice Mean ± SEM; *p<0.05; n=6 or 7; unpaired two-tailed Student’s *t*-test.

### AP39 increased UCP1 expression in VSMCs in atherosclerotic lesions

To further confirm upregulated expression of UCP1 upon administration of mitochondrial H_2_S donor AP39, we investigated alterations in UCP1 protein expression in the most abundant cells in atherosclerotic plaque, VSMCs. Indeed, immunohistochemical staining of atherosclerotic lesions revealed increased abundance of cells positive for both VSMC marker – calponin and UCP1 upon treatment with AP39 (Fig.6A and B, Supplemental Figure 6).

**Figure 6.**
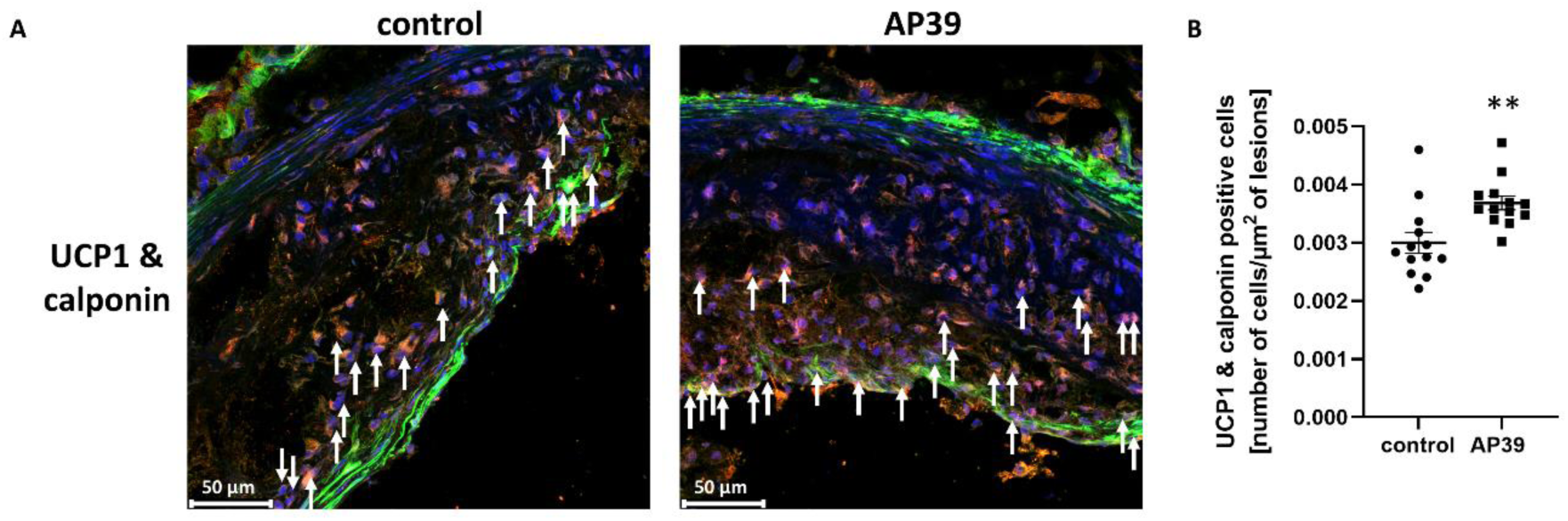
AP39 increases UCP1 expression in VSMCs in atherosclerotic plaque. Representative immunohistochemical staining of aortic roots showing calponin (green), DAPI (blue) and UCP1 (red) co-localization in control and AP39-treated mice **(A)**. White arrows indicate some of the UCP1 & calponin positive cells. Quantitative analysis of UCP1 expression in VSMCs in atherosclerotic lesions **(B)**. Mean ± SEM; **p<0.01 compared to control mice; n=13; unpaired two-tailed Student’s *t*-test.

### AP39 increased eNOS expression and activation in the aorta of apoE^-/-^ mice

Our LC‒MS/MS analysis also showed the increased expression of eNOS at the protein level in the aorta of apoE^-/-^ mice (Supplemental Table 1). Similarly, RT-qPCR analysis indicated upregulated mRNA expression for *Nos3* (eNOS) (Fig. 7D) in the aorta of apoE^-/-^ mice. The main function of eNOS is the production of NO, therefore, to assess eNOS activity, we measured its phosphorylation on serine 1177 by Western blot. Indeed, AP39 administration increased phosphorylation of eNOS on serine 1177 normalized to total eNOS level (Fig. 7A and B), thus impacting on eNOS activity and NO production. However, eNOS expression at the protein level measured by Western blot was not statistically significant (Fig. 7A and C). Uncropped Western blot images are presented in Supplemental Figure 4.

**Figure 7.**
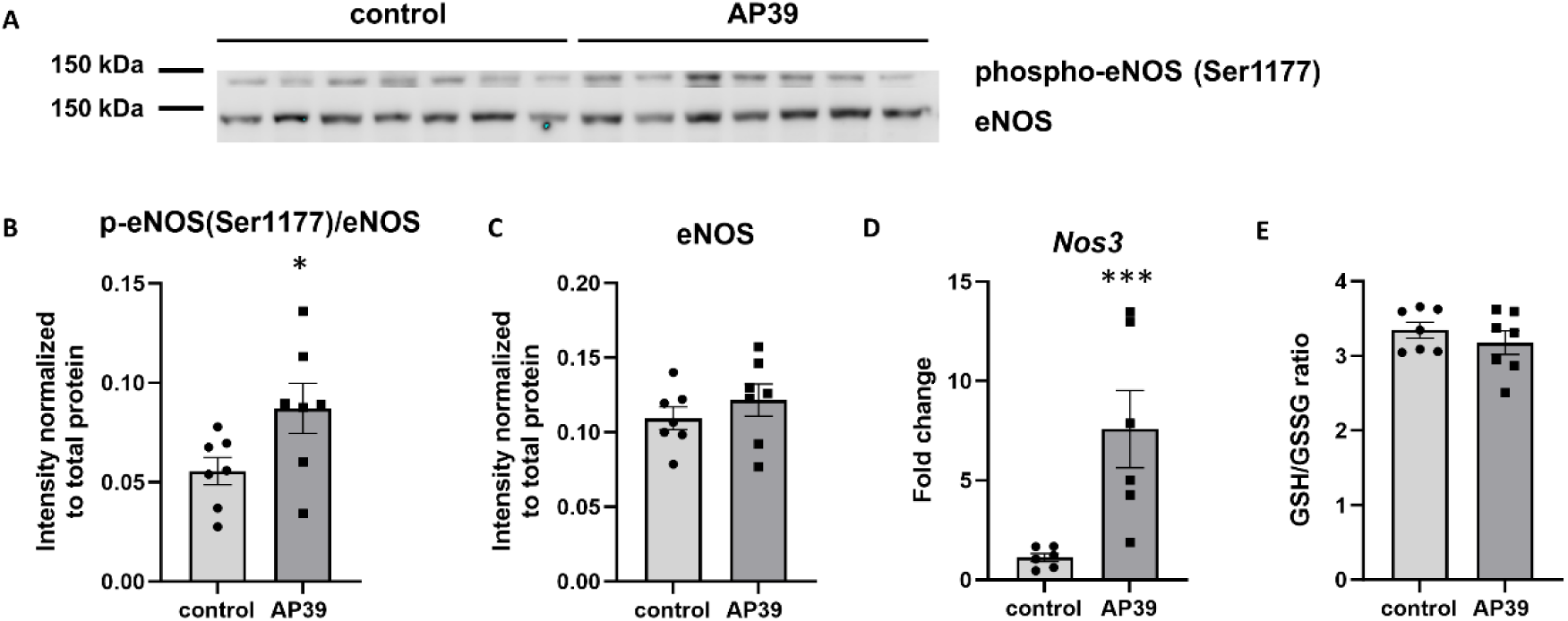
AP39 increases eNOS activation in the aorta of apoE^-/-^ mice. Western blot of p-eNOS (Ser1177) and eNOS in the aorta of control and AP39-treated mice **(A)**. Quantitative analysis of Western blots for p-eNOS(Ser1177)/eNOS **(B)** and eNOS **(C)** expression in the aorta of control and AP39-treated mice. *Nos3* mRNA expression in the aorta of control and AP39-treated mice **(D)**. The ratio of GSH to GSSG in the plasma of control and AP39-treated mice (**E**). Mean ± SEM; *p<0.05, ***p<0.001; n=6 or 7; unpaired two-tailed Student’s *t*-test.

To assess oxidative stress level upon AP39 administration in apoE^-/-^ mice, we measured GSH/GSSG ratio, which is an indicator of cellular health and a marker of oxidative stress. Our results indicated that GSH/GSSG ratio did not change because of AP39 treatment in the plasma of apoE^-/-^ mice (Fig. 7E).

## Discussion

A growing body of evidence points to the role of mitochondrial dysfunction in the pathogenesis of atherosclerosis. In the present study, we applied mitochondrial H_2_S donor AP39, which has potential to stimulate cellular bioenergetics and improve mitochondrial function, to evaluate atherosclerosis development in apoE^-/-^ mice. We demonstrated for the first time that AP39 **1)** reduced and stabilized atherosclerotic plaque, **2)** decreased proinflammatory M1 macrophages and increased anti-inflammatory M2 macrophages in atherosclerotic lesions, **3)** upregulated pathways related to mitochondrial function, such as cellular respiration and fatty acid β-oxidation, **4)** increased UCP1 protein expression in VSMCs in atherosclerotic lesions, **5)** and upregulated mRNA expression of other thermogenesis-related genes in the aorta but not PVAT of apoE^-/-^ mice.

Our results indicated that mitochondria-targeted, slow-releasing H_2_S donor AP39 administrated at a dose (0.1mg/kg/day) for 16 weeks reduced atherosclerosis in apoE^-/-^ mice. Moreover, AP39 stabilized atherosclerotic plaque as shown by decreased macrophage content and increased collagen depositions in atherosclerotic lesions of apoE^-/-^ mice. However, it did not influence the area of necrotic core and SMA content. This is in line with other studies showing the importance of H_2_S in the pathogenesis of atherosclerosis. It has been observed that the deficiency of H_2_S caused by the knockout of CSE enzyme accelerated atherosclerosis in apoE^-/-^ mice ^16^. In addition, administration of exogenous H_2_S in the form of inorganic salts (NaSH) ^17^ or slow-releasing H_2_S donor (GYY4137) ^18^ was shown to attenuate atherosclerosis in apoE^-/-^ mice. Among others, the beneficial actions of H_2_S in atherosclerosis reduction were related to PI3K/Akt and TLR4 signaling ^18^, inhibition of NLRP3 inflammasome ^19^, upregulation of protein S-nitrosylation ^20^ and activation of Nrf2/HO-1 signaling pathway ^21^. However, in the present study, we focused on investigation the mechanisms of action of mitochondria-targeted H_2_S donor AP39 in atherosclerosis development with the regard to mitochondria-related pathways.

Decreased macrophage content in atherosclerotic lesions upon AP39 administration prompted us to explore plaque composition more deeply in terms of macrophage phenotypes. We showed that AP39 treatment reduced the level of proinflammatory M1 macrophages and elevated the level of anti-inflammatory M2 macrophages in atherosclerotic lesions of apoE^-/-^ mice. The ability of H_2_S to influence macrophage polarization was also confirmed in another study showing that H_2_S donor NaSH induced M2 polarization in myocardial infarction ^22^. Generally, two main phenotypes of macrophages: proinflammatory M1 macrophages and anti-inflammatory M2 macrophages, that exhibit different mitochondrial metabolism, can be distinguished in atherosclerotic plaques. The M1 macrophages rely on glycolysis and are responsible for the clearance of pathogens, whereas M2 macrophages exploit oxidative phosphorylation as a main energy source and play a role in the resolution of inflammation, tissue repair and wound healing ^5^. Importantly, it has been shown that macrophage phenotype may have an impact on plaque vulnerability. More M1 macrophages were observed in unstable plaque and more M2 macrophages were seen in stable plaque in patients with acute ischemic attack ^23^. In addition, several studies have demonstrated that M1 macrophage phenotype is linked to atherosclerosis progression ^24^ and in turn, the induction of macrophage polarization to M2 by IL-13 could reduce disease progression ^25^. M2 macrophages can promote the removal of dead cells by efferocytosis and tissue repair through collagen production, thus they may prevent the progression of atherosclerosis or even cause plaque regression ^26^. Therefore, macrophage phenotype switching from proinflammatory M1 to anti-inflammatory M2 type observed upon treatment with mitochondrial donor of H_2_S AP39 may represent a promising strategy in the treatment of atherosclerosis. However, whether AP39 could switch macrophage polarization to M2 phenotype through the reprogramming of mitochondrial metabolism in macrophages, requires further investigations.

Importantly, our proteomic data revealed downregulated expression of some proteins regulating macrophage polarization, such as DPPI, DPP9, GMFG and LCN2 in the aorta of AP39-treated apoE^-/-^ mice. It has been shown that upregulated DPPI, a lysosomal cysteine protease, induced polarization of macrophages to proinflammatory M1 phenotype through FAK-triggered p38 MAPK/NF-κβ pathway ^27^, whereas its deficiency attenuated atherogenesis and led to anti-inflammatory M2 polarization ^28^. Similarly, inhibition of DPP8/9, serine proteolytic enzymes that are expressed in monocytes and macrophages, decreased M1 macrophage polarization and reduced atherosclerosis ^29^. On the other hand, it has been found that GMFG is a regulator of cellular iron metabolism and macrophage phenotype and its knockdown promoted M2 macrophage polarization through mitochondrial ROS formation ^30^. Importantly, LCN2, a bioactive hormone expressed in macrophages, adipose tissue, and neutrophils, was shown to exert pro-atherosclerotic effects by increasing M1 macrophage polarization, which was associated with NF-κβ upregulation ^31^ and NLRP3 inflammasome activation ^32^. To sum up, downregulated expression of DPPI, DPP9 and LCN2 could lead to decreased proinflammatory M1 macrophage activation, whereas reduced expression of GMFG may promote anti-inflammatory M2 macrophage polarization. Whether these alterations in protein expression translate into decreased proinflammatory M1 macrophages and increased anti-inflammatory M2 macrophages in atherosclerotic lesions upon AP39 administration, require further investigations.

Notably, our DIA quantitative proteomic approach indicated increased expression of different collagen types (e.g., COL1A1, COL5A1) and decreased expression of MMPs (MMP-2, -3 -12 and -19) as well as macrophage and monocyte markers (CD68, CD14, MPEG1) in the aorta of AP39-treated mice. These results are in line with the immunohistochemistry data and may confirm increased stabilization of atherosclerotic plaque upon AP39 administration. Of note, MMP-3 is involved in the degradation of the fibrous cap ^33^, whereas MMP-12 may be critical to the progression of atherosclerosis via degradation of the basement membrane ^34^. Interestingly, deficiency of MMP-2 was shown to reduce atherosclerotic lesions in apoE^-/-^ mice ^35^. Thus, decreasing the expression of MMPs by AP39 may have a positive impact on atherosclerosis development. Apart from that, our proteomic analysis also pointed out to the downregulation of pathways related to immune system, platelet activation and aggregation, and complement and coagulation cascades upon AP39 treatment in the aorta of apoE^-/-^ mice. Particularly, expression of TXA was reduced in the aorta of AP39-treated mice. It has been shown that the inhibition of thromboxane A2 synthesis or knockout of the thromboxane A2 receptor delayed atherogenesis in murine models ^36^. Moreover, anti-inflammatory and anti-thrombotic effects of H_2_S have been well-described ^37^, and our results also confirmed this for mitochondria-targeted H_2_S donor AP39. Importantly, decreased expression of proteins engaged in inflammation and thrombosis in the aorta may contribute to the beneficial action of AP39 in the reduction of atherosclerosis.

Interestingly, our MS data showed increased expression of proteins involved in cellular respiration: oxidative phosphorylation and Krebs cycle as well as mitochondrial fatty acid β-oxidation upon administration of mitochondrial H_2_S donor AP39. We observed upregulated expression of all enzymes of ETC (Complex I – IV) and most of the enzymes of Krebs cycle (e.g., CS, ACO2, PDHB) and β-oxidation of fatty acids (e.g., ACAA2, SCAD, HADHA/B) in the aorta of AP39-treated mice. These results point to the mitochondrial action of AP39 in the aorta of apoE^-/-^ mice and may indicate improved mitochondrial function and energy production. Furthermore, bioinformatic analysis of proteomic data also revealed upregulated thermogenesis pathway in the aorta of AP39-treated mice. Notably, protein expression of UCP1, HSL, ABHD5 and CPT2 increased upon AP39 treatment in the aorta. In addition, we showed increased mRNA expression for transcription factors and regulators involved in thermogenesis, such as *Cidea*, *Pgc1*, *Pparg* and *Nrf2* in the aorta of AP39-treated mice. Importantly, upregulated thermogenesis pathway was observed only in the aorta but not PVAT of apoE^-/-^ mice upon AP39 administration. UCP1 is a mitochondrial protein that dissipates the mitochondrial electrochemical gradient in the form of heat rather than energy production. It is specifically expressed in brown and beige adipocytes; however, it can be induced in white adipocytes in response to various stimuli, including cold exposure, insulin and β3 agonists, which is called the browning process ^38^. Transcription of UCP1 is regulated by several transcription factors, including *Pgc1*, *Pparg*, *Cidea* and *Nrf2*. *Pgc1* and *Pparg* transcription factors form complex and bind to the promotor of *Ucp1* gene. *Pgc1* is not only a crucial regulator in thermogenesis but also in the mitochondrial biogenesis ^39^. *Cidea* is a transcriptional regulator that is necessary to induce UCP1 expression and browning of adipose tissue ^40^. Similarly, *Nrf2* transcription factor was shown to induce UCP1 expression in adipocytes ^41^. Thermogenic activity of brown and beige adipocytes is driven by free fatty acids that enter the mitochondria where they are subjected to β-oxidation and catabolic processing in Krebs cycle to fuel the ETC. ETC generates a proton gradient across the inner mitochondrial membrane that is disrupted by UCP1 ^42^. The entry of free fatty acids to mitochondria is possible thanks to the activity of HSL and ABHD5 enzymes that are engaged in hydrolysis of fatty acids from triacylglycerol as well as the activity of CPT2 enzyme that transports long chain fatty acids to mitochondria for β-oxidation ^43^. Noteworthy, our proteomic analysis pointed out to the increased expression of UCP1, HSL, ABHD5, CPT2 as well as proteins involved in β-oxidation, Krebs cycle and oxidative phosphorylation in the aorta of AP39-treated mice.

It should be emphasized that our proteomic and mRNA data strongly supported upregulated thermogenesis pathway in the aorta but not PVAT of AP39-treated mice. PVAT resembles both brown and white adipose tissues. The expression of UCP1 is thought to be reserved for brown and beige adipose tissues, however it was also detected in the aorta ^44^. Interestingly, it has been shown that UCP1 deficiency in PVAT exacerbates vascular inflammation, endothelial dysfunction and atherosclerosis through the activation of NLRP3 inflammasome and IL-1β production ^45^, pointing to a protective role of UCP1 in atherogenesis. Moreover, polarization of macrophages to M2 phenotype was shown to induce UCP1 expression and thermogenesis leading to browning of white adipose tissue ^46^. Our results indicated that mitochondrial H_2_S could increase both M2 polarization and thermogenesis. Of note, there is little evidence about the role of H_2_S in thermogenesis. One study has shown that endogenous peripheral H_2_S increased thermogenesis in brown adipose tissue in cold environments in rats ^47^. Importantly, we observed increased thermogenesis upon AP39 administration in the aorta of mice, not classical adipose tissue, such as PVAT that surrounds the aorta. It is tempting to speculate that upregulated thermogenesis in the aorta is limited to VSMCs, which are the most abundant cells in the aorta and atherosclerotic lesions. Indeed, our results showed increased expression of UCP1 in VSMCs in atherosclerotic lesions upon AP39 administration. VSMCs are very plastic and can dedifferentiate to different phenotypes, including adipocyte-like VSMCs that are classified as beige adipocytes and express UCP1 protein ^48^. The relevance of this phenotype in atherosclerotic lesions is not known, but presumably adipocyte-like VSMCs may burn fatty acids and produce heat in the presence of UCP1 leading to reduced lipid content in atherosclerotic plaques. However, further studies are needed to confirm this attractive hypothesis regarding the role of mitochondrial H_2_S in inducing thermogenesis in adipocyte-like VSMCs in atherosclerotic lesions as a potential therapeutic target.

Finally, we found that mitochondrial donor of H_2_S AP39 increased eNOS expression as well as eNOS activation measured as eNOS phosphorylation on serine 1177 in the aorta of apoE^-/-^ mice. Our results are in line with other studies that showed H_2_S role in modulating the activity of eNOS and hence increased NO production. It has been found that H_2_S activates PI3K/Akt/eNOS pathway in endothelial cells ^49^ and directly interacts with eNOS by sulfhydration of critical cysteines, which stabilizes eNOS in dimeric state and optimizes eNOS-derived NO production ^50^. Of note, NO is a potent vasodilator and anti-inflammatory agent, and its reduced bioavailability leads to endothelial dysfunction associated with the development of atherosclerosis ^51^. Our proteomic data also revealed that administration of mitochondrial H_2_S donor AP39 decreased expression of proteins characteristic of activated endothelium, such as P-selectin and VCAM-1 in the aorta of apoE^-/-^ mice, which could indicate improved endothelial function. Moreover, upon AP39 treatment in the aorta, we found downregulated different subunits of neutrophilic type NADPH oxidase, which serves as the main source of ROS in vessels during atherogenesis ^52^. However, we did not observe altered GSH/GSSG ratio in the plasma of AP39-treated mice, which is an indicator of oxidative stress level. Overall, AP39 administration may improve endothelial function by increasing NO production and reducing ROS level, which may have a beneficial impact in the treatment of atherosclerosis.

Our study has some limitations. We elucidated the mechanism of action of mitochondrial donor of H_2_S AP39 in atherosclerosis using quantitative DIA proteomics in the aorta of apoE^-/-^ mice. It would be more relevant to perform proteomic analysis solely in atherosclerotic lesions, however it requires more sophisticated technology, such as combination of laser capture microdissection and single-cell proteomics, methods that are still being under development/improvement.

Overall, our results indicated that mitochondrial H_2_S donor AP39 reduced atherosclerosis by reprogramming macrophages from proinflammatory M1 to anti-inflammatory M2 phenotype in atherosclerotic plaque and upregulating cellular respiration, mitochondrial fatty acids β-oxidation and thermogenesis in the aorta of apoE^-/-^ mice. Moreover, it increased the expression of UCP1 in VSMCs in atherosclerotic lesions. It is tempting to speculate that mitochondrial donor of H_2_S AP39 could provide potentially a novel therapeutic approach to the treatment/prevention of atherosclerosis.

## Conclusions

Our findings demonstrated that mitochondrial donor of H_2_S AP39 reduced atherosclerosis and stabilized atherosclerotic lesions in apoE^-/-^ mice by decreasing macrophage content and increasing collagen depositions within lesions. Moreover, AP39 administration reprogrammed macrophages from proinflammatory M1 to anti-inflammatory M2 in atherosclerotic plaques. Importantly, AP39 downregulated pathways associated with immune system, platelet activation and aggregation, complement and coagulation cascades, and upregulated pathways related to mitochondrial function, such as cellular respiration, fatty acid β-oxidation and thermogenesis in the aorta, which may point out to the improved mitochondrial function. Furthermore, treatment with AP39 increased the expression of UCP1 and other proteins related to thermogenesis in the aorta but not in PVAT of apoE^-/-^ mice. Immunohistochemical staining showed that UCP1 expression was increased in VSMCs in atherosclerotic lesions upon AP39 administration. These results suggested that mitochondria-targeted H_2_S could be a novel therapeutic strategy in the treatment of atherosclerosis. We hope that knowledge regarding the mechanisms of action of mitochondrial H_2_S can provide new insights for the pathogenesis of atherosclerosis and facilitate the advancement of new therapeutic targets.

**Figure 8.**
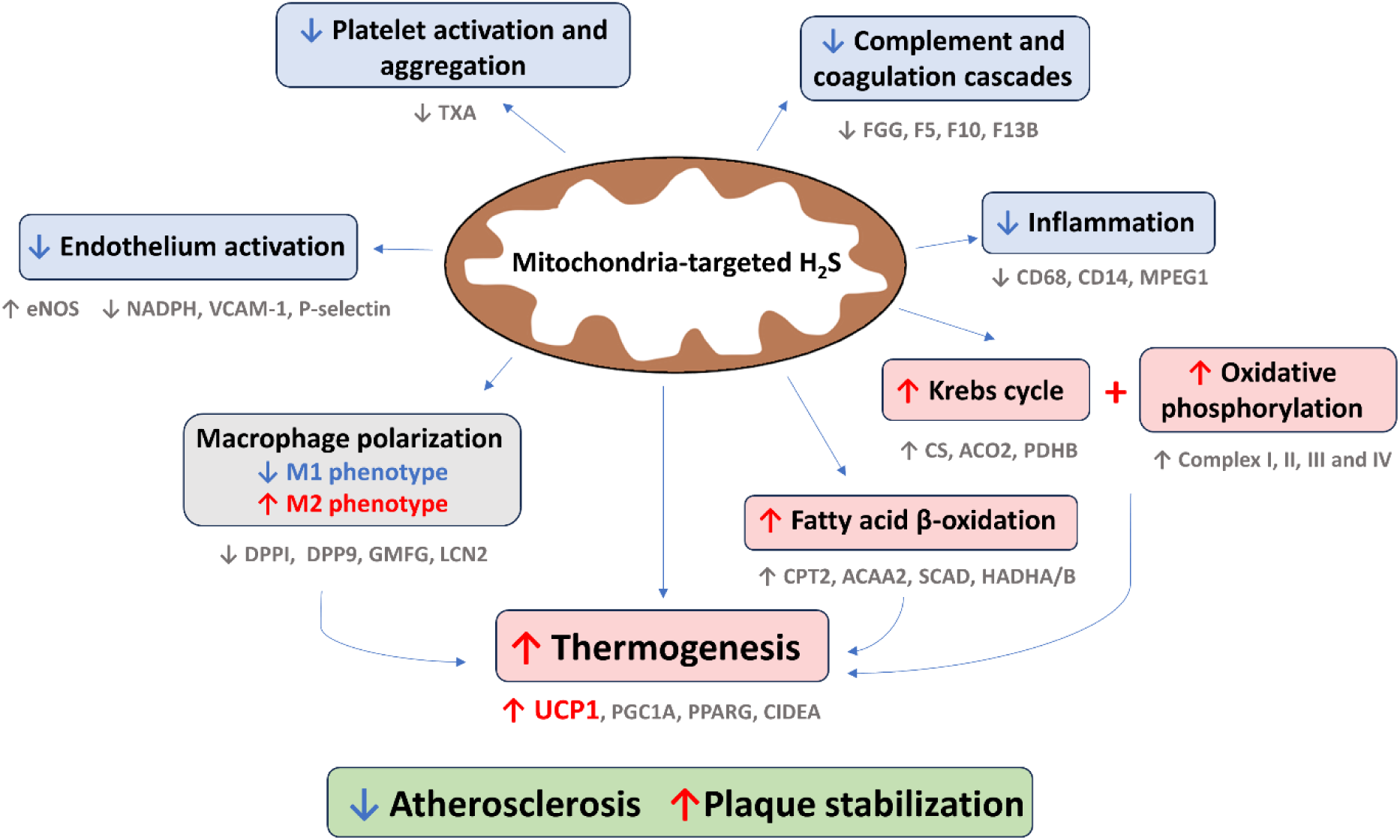
The summary of beneficial effects of mitochondria-targeted H2S donor AP39 on atherosclerosis development.

## Abbreviations

3-MST: 3-mercaptopyruvate sulfurtransferase
ABHD5: 1-acylglycerol-3-phosphate O-acyltransferase ABHD5
ACAA2: 3-ketoacyl-CoA thiolase
ACO2: aconitate hydratase
ANG-2: angiopoietin-2
CBS: cystathionine-β-synthase
CD14: monocyte differentiation antigen
CD14: Cidea cell death inducing
DFFA: like effector A
COL1A1: collagen alpha-1(I) chain
COX IV: cytochrome c oxidase subunit 4 isoform 1
CPT2: carnitine O-palmitoyltransferase 2
CS: citrate synthase
CSE: cystathionine gamma-lyase
DDA: data-dependent acquisition
DIA: data-independent acquisition
DPP9: dipeptidyl peptidase 9
DPPI: dipeptidyl peptidase 1
Elovl3: ELOVL fatty acid elongase 3
eNOS: Nos3 endothelial nitric oxide synthase
ETC: electron transport chain
FC: fold change
GMFG: glia maturation factor gamma
GSH: reduced glutathione
GSSG: oxidized glutathione
HADHA/B: trifunctional enzyme subunit alpha/beta
HFD: high fat diet
HSL: hormone-sensitive lipase
ICAM: intercellular adhesion molecule 1
IDH3A: isocitrate dehydrogenase [NAD] subunit alpha
IDH3G: isocitrate dehydrogenase [NAD] subunit gamma 1
iNOS: nitric oxide synthase 2
LC‒MS/MS: liquid chromatography – tandem mass spectrometry
LCN2: lipocalin-2
M-CSF: macrophage colony-stimulating factor
MMP: matrix metalloproteinase
MPEG1: macrophage-expressed gene 1 protein
Nrf2: nuclear factor erythroid 2-related factor 2
PAI-1: plasminogen activator inhibitor-1
PDHB: pyruvate dehydrogenase E1 component subunit beta
Pgc1: peroxisome proliferator-activated receptor gamma coactivator 1-alpha
PINE: protein interaction network extractor
Pparg: peroxisome proliferator activated receptor gamma
PVAT: perivascular adipose tissue
SCAD: short-chain specific acyl-CoA dehydrogenase
SDHA: succinate dehydrogenase [ubiquinone] flavoprotein subunit
SDHB: succinate dehydrogenase iron-sulfur subunit
SMA: smooth muscle α-actin
TXA: thromboxane-A synthase
UCP1: mitochondrial brown fat uncoupling protein 1
VCAM-1: vascular cell adhesion protein 1
VEGF: vascular endothelial growth factor
VSMC: vascular smooth muscle cell

## Author contributions

AS was responsible for the conception and design of the study. AS, AW, KC, BP, AS, MS, BKC, MEW, RT, MW and KK were responsible for analyses of the samples. AS and RO were responsible for the interpretation of data. AS drafted the article.

## References

1. Virmani R, Kolodgie FD, Burke AP, Farb A, Schwartz SM. Lessons from sudden coronary death: a comprehensive morphological classification scheme for atherosclerotic lesions. Arterioscler Thromb Vasc Biol 2000;20:1262–1275.

2. Zhou R, Yazdi AS, Menu P, Tschopp J. A role for mitochondria in NLRP3 inflammasome activation. Nature 2011;469:221–225.

3. Peng W, Cai G, Xia Y, Chen J, Wu P, Wang Z, Li G, Wei D. Mitochondrial Dysfunction in Atherosclerosis. DNA Cell Biol 2019;38:597–606.

4. Mills EL, O’Neill LA. Reprogramming mitochondrial metabolism in macrophages as an anti-inflammatory signal. Eur J Immunol 2016;46:13–21.

5. Mosser DM, Edwards JP. Exploring the full spectrum of macrophage activation. Nat Rev Immunol 2008;8:958–969.

6. Wang Z-J, Wu J, Guo W, Zhu Y-Z. Atherosclerosis and the Hydrogen Sulfide Signaling Pathway - Therapeutic Approaches to Disease Prevention. Cell Physiol Biochem Int J Exp Cell Physiol Biochem Pharmacol 2017;42:859–875.

7. Kabil O, Motl N, Banerjee R. H2S and its role in redox signaling. Biochim Biophys Acta 2014;1844:1355– 1366.

8. Nicholls P. Inhibition of cytochrome c oxidase by sulphide. Biochem Soc Trans 1975;3:316–319.

9. Módis K, Coletta C, Erdélyi K, Papapetropoulos A, Szabo C. Intramitochondrial hydrogen sulfide production by 3-mercaptopyruvate sulfurtransferase maintains mitochondrial electron flow and supports cellular bioenergetics. FASEB J Off Publ Fed Am Soc Exp Biol 2013;27:601–611.

10. Szczesny B, Módis K, Yanagi K, Coletta C, Le Trionnaire S, Perry A, Wood ME, Whiteman M, Szabo C. AP39, a novel mitochondria-targeted hydrogen sulfide donor, stimulates cellular bioenergetics, exerts cytoprotective effects and protects against the loss of mitochondrial DNA integrity in oxidatively stressed endothelial cells in vitro. Nitric Oxide Biol Chem 2014;41:120–130.

11. Stachowicz A, Wiśniewska A, Kuś K, Białas M, Łomnicka M, Totoń-Żurańska J, Kiepura A, Stachyra K, Suski M, Bujak-Giżycka B, Jawień J, Olszanecki R. Diminazene Aceturate Stabilizes Atherosclerotic Plaque and Attenuates Hepatic Steatosis in apoE-Knockout Mice by Influencing Macrophages Polarization and Taurine Biosynthesis. Int J Mol Sci 2021;22:5861.

12. Trionnaire SL, Perry A, Szczesny B, Szabo C, Winyard PG, Whatmore JL, Wood ME, Whiteman M. The synthesis and functional evaluation of a mitochondria-targeted hydrogen sulfide donor, (10-oxo-10-(4-(3-thioxo-3H-1,2-dithiol-5-yl)phenoxy)decyl)triphenylphosphonium bromide (AP39). MedChemComm 2014;5:728–736.

13. Magierowska K, Wójcik-Grzybek D, Korbut E, Bakalarz D, Ginter G, Danielak A, Kwiecień S, Chmura A, Torregrossa R, Whiteman M, Magierowski M. The mitochondria-targeted sulfide delivery molecule attenuates drugs-induced gastropathy. Involvement of heme oxygenase pathway. Redox Biol 2023;66:102847.

14. Sundararaman N, Go J, Robinson AE, Mato JM, Lu SC, Van Eyk JE, Venkatraman V. PINE: An Automation Tool to Extract and Visualize Protein-Centric Functional Networks. J Am Soc Mass Spectrom 2020;31:1410–1421.

15. Vizcaíno JA, Csordas A, Del-Toro N, Dianes JA, Griss J, Lavidas I, Mayer G, Perez-Riverol Y, Reisinger F, Ternent T, Xu Q-W, Wang R, Hermjakob H. 2016 update of the PRIDE database and its related tools. Nucleic Acids Res 2016;**44**:D447–456.

16. Mani S, Li H, Untereiner A, Wu L, Yang G, Austin RC, Dickhout JG, Lhoták Š, Meng QH, Wang R. Decreased endogenous production of hydrogen sulfide accelerates atherosclerosis. Circulation 2013;127:2523–2534.

17. Xiong Q, Wang Z, Yu Y, Wen Y, Suguro R, Mao Y, Zhu YZ. Hydrogen sulfide stabilizes atherosclerotic plaques in apolipoprotein E knockout mice. Pharmacol Res 2019;144:90–98.

18. Zheng Y, Lv P, Huang J, Ke J, Yan J. GYY4137 exhibits anti-atherosclerosis effect in apolipoprotein E (-/-) mice via PI3K/Akt and TLR4 signalling. Clin Exp Pharmacol Physiol 2020;**47**:1231–1239.

19. Yue L-M, Gao Y-M, Han B-H. Evaluation on the effect of hydrogen sulfide on the NLRP3 signaling pathway and its involvement in the pathogenesis of atherosclerosis. J Cell Biochem 2019;120:481–492.

20. Lin Y, Chen Y, Zhu N, Zhao S, Fan J, Liu E. Hydrogen sulfide inhibits development of atherosclerosis through up-regulating protein S-nitrosylation. Biomed Pharmacother Biomedecine Pharmacother 2016;83:466–476.

21. Xie L, Gu Y, Wen M, Zhao S, Wang W, Ma Y, Meng G, Han Y, Wang Y, Liu G, Moore PK, Wang X, Wang H, Zhang Z, Yu Y, Ferro A, Huang Z, Ji Y. Hydrogen Sulfide Induces Keap1 S-sulfhydration and Suppresses Diabetes-Accelerated Atherosclerosis via Nrf2 Activation. Diabetes 2016;65:3171–3184.

22. Miao L, Shen X, Whiteman M, Xin H, Shen Y, Xin X, Moore PK, Zhu Y-Z. Hydrogen Sulfide Mitigates Myocardial Infarction via Promotion of Mitochondrial Biogenesis-Dependent M2 Polarization of Macrophages. Antioxid Redox Signal 2016;25:268–281.

23. Cho KY, Miyoshi H, Kuroda S, Yasuda H, Kamiyama K, Nakagawara J, Takigami M, Kondo T, Atsumi T. The phenotype of infiltrating macrophages influences arteriosclerotic plaque vulnerability in the carotid artery. J Stroke Cerebrovasc Dis Off J Natl Stroke Assoc 2013;22:910–918.

24. Hanna RN, Shaked I, Hubbeling HG, Punt JA, Wu R, Herrley E, Zaugg C, Pei H, Geissmann F, Ley K, Hedrick CC. NR4A1 (Nur77) deletion polarizes macrophages toward an inflammatory phenotype and increases atherosclerosis. Circ Res 2012;110:416–427.

25. Cardilo-Reis L, Gruber S, Schreier SM, Drechsler M, Papac-Milicevic N, Weber C, Wagner O, Stangl H, Soehnlein O, Binder CJ. Interleukin-13 protects from atherosclerosis and modulates plaque composition by skewing the macrophage phenotype. EMBO Mol Med 2012;4:1072–1086.

26. Bi Y, Chen J, Hu F, Liu J, Li M, Zhao L. M2 Macrophages as a Potential Target for Antiatherosclerosis Treatment. Neural Plast 2019;2019:6724903.

27. Alam S, Liu Q, Liu S, Liu Y, Zhang Y, Yang X, Liu G, Fan K, Ma J. Up-regulated cathepsin C induces macrophage M1 polarization through FAK-triggered p38 MAPK/NF-κB pathway. Exp Cell Res 2019;382:111472.

28. Herías V, Biessen EAL, Beckers C, Delsing D, Liao M, Daemen MJ, Pham CCTN, Heeneman S. Leukocyte cathepsin C deficiency attenuates atherosclerotic lesion progression by selective tuning of innate and adaptive immune responses. Arterioscler Thromb Vasc Biol 2015;35:79–86.

29. Waumans Y, Vliegen G, Maes L, Rombouts M, Declerck K, Van Der Veken P, Vanden Berghe W, De Meyer GRY, Schrijvers D, De Meester I. The Dipeptidyl Peptidases 4, 8, and 9 in Mouse Monocytes and Macrophages: DPP8/9 Inhibition Attenuates M1 Macrophage Activation in Mice. Inflammation 2016;39:413–424.

30. Aerbajinai W, Ghosh MC, Liu J, Kumkhaek C, Zhu J, Chin K, Rouault TA, Rodgers GP. Glia maturation factor-γ regulates murine macrophage iron metabolism and M2 polarization through mitochondrial ROS. Blood Adv 2019;3:1211–1225.

31. Shibata K, Sato K, Shirai R, Seki T, Okano T, Yamashita T, Koide A, Mitsuboshi M, Mori Y, Hirano T, Watanabe T. Lipocalin-2 exerts pro-atherosclerotic effects as evidenced by in vitro and in vivo experiments. Heart Vessels 2020;35:1012–1024.

32. Kim SL, Shin MW, Kim SW. Lipocalin 2 activates the NLRP3 inflammasome via LPS-induced NF-κB signaling and plays a role as a pro-inflammatory regulator in murine macrophages. Mol Med Rep 2022;26:358.

33. Magdalena K, Magdalena K, Grazyna S. The Role of Matrix Metalloproteinase-3 In the Development of Atherosclerosis and Cardiovascular Events. EJIFCC 2006;17:2–5.

34. Yamada S, Wang K-Y, Tanimoto A, Fan J, Shimajiri S, Kitajima S, Morimoto M, Tsutsui M, Watanabe T, Yasumoto K, Sasaguri Y. Matrix Metalloproteinase 12 Accelerates the Initiation of Atherosclerosis and Stimulates the Progression of Fatty Streaks to Fibrous Plaques in Transgenic Rabbits. Am J Pathol 2008;172:1419–1429.

35. Kuzuya M, Nakamura K, Sasaki T, Wu Cheng X, Itohara S, Iguchi A. Effect of MMP-2 Deficiency on Atherosclerotic Lesion Formation in ApoE-Deficient Mice. Arterioscler Thromb Vasc Biol 2006;26:1120– 1125.

36. Cyrus T, Yao Y, Ding T, Dogné JM, Praticò D. Thromboxane receptor blockade improves the antiatherogenic effect of thromboxane A2 suppression in LDLR KO mice. Blood 2007;109:3291–3296.

37. 37. Gemici B, Wallace JL. Chapter Nine - Anti-inflammatory and Cytoprotective Properties of Hydrogen Sulfide. In: Cadenas E, Packer L, eds. Methods in Enzymology. Academic Press; 2015. p169–193.

38. Harms M, Seale P. Brown and beige fat: development, function and therapeutic potential. Nat Med 2013;19:1252–1263.

39. Puigserver P, Wu Z, Park CW, Graves R, Wright M, Spiegelman BM. A cold-inducible coactivator of nuclear receptors linked to adaptive thermogenesis. Cell 1998;92:829–839.

40. Jash S, Banerjee S, Lee M-J, Farmer SR, Puri V. CIDEA Transcriptionally Regulates UCP1 for Britening and Thermogenesis in Human Fat Cells. iScience 2019;20:73–89.

41. Chang S-H, Jang J, Oh S, Yoon J-H, Jo D-G, Park UJY& KW. Nrf2 induces Ucp1 expression in adipocytes in response to β3-AR stimulation and enhances oxygen consumption in high-fat diet-fed obese mice. BMB Rep 2021;54:419–424.

42. Roth CL, Molica F, Kwak BR. Browning of White Adipose Tissue as a Therapeutic Tool in the Fight against Atherosclerosis. Metabolites 2021;11:319.

43. Ying Z, Tramper N, Zhou E, Boon MR, Rensen PCN, Kooijman S. Role of thermogenic adipose tissue in lipid metabolism and atherosclerotic cardiovascular disease: lessons from studies in mice and humans. Cardiovasc Res 2022;119:905–918.

44. Winn NC, Grunewald ZI, Gastecki ML, Woodford ML, Welly RJ, Clookey SL, Ball JR, Gaines TL, Karasseva NG, Kanaley JA, Sacks HS, Vieira-Potter VJ, Padilla J. Deletion of UCP1 enhances ex vivo aortic vasomotor function in female but not male mice despite similar susceptibility to metabolic dysfunction. Am J Physiol - Endocrinol Metab 2017;313:E402–E412.

45. Gu P, Hui X, Zheng Q, Gao Y, Jin L, Jiang W, Zhou C, Liu T, Huang Y, Liu Q, Nie T, Wang Y, Wang Y, Zhao J, Xu A. Mitochondrial uncoupling protein 1 antagonizes atherosclerosis by blocking NLRP3 inflammasome-dependent interleukin-1β production. Sci Adv 2021;7:eabl4024.

46. Qiu Y, Nguyen KD, Odegaard JI, Cui X, Tian X, Locksley RM, Palmiter RD, Chawla A. Eosinophils and type 2 cytokine signaling in macrophages orchestrate development of functional beige fat. Cell 2014;157:1292–1308.

47. Soriano RN, Braga SP, Breder JSC, Batalhao ME, Oliveira-Pelegrin GR, Ferreira LFR, Rocha MJA, Carnio EC, Branco LGS. Endogenous peripheral hydrogen sulfide is propyretic: its permissive role in brown adipose tissue thermogenesis in rats. Exp Physiol 2018;103:397–407.

48. Long JZ, Svensson KJ, Tsai L, Zeng X, Roh HC, Kong X, Rao RR, Lou J, Lokurkar I, Baur W, Castellot JJ, Rosen ED, Spiegelman BM. A smooth muscle-like origin for beige adipocytes. Cell Metab 2014;19:810– 820.

49. Lin F, Yang Y, Wei S, Huang X, Peng Z, Ke X, Zeng Z, Song Y. Hydrogen Sulfide Protects Against High Glucose-Induced Human Umbilical Vein Endothelial Cell Injury Through Activating PI3K/Akt/eNOS Pathway. Drug Des Devel Ther 2020;14:621–633.

50. Szabo C. Hydrogen sulfide, an enhancer of vascular nitric oxide signaling: mechanisms and implications. Am J Physiol-Cell Physiol 2017;312:C3–C15.

51. Kawashima S, Yokoyama M. Dysfunction of Endothelial Nitric Oxide Synthase and Atherosclerosis. Arterioscler Thromb Vasc Biol 2004;24:998–1005.

52. Poznyak AV, Grechko AV, Orekhova VA, Khotina V, Ivanova EA, Orekhov AN. NADPH Oxidases and Their Role in Atherosclerosis. Biomedicines 2020;8:206.

