## Supplemental Figures for "Mitochondria-targeted hydrogen sulfide donor reduces atherogenesis by reprogramming macrophages and increasing UCP1 expression in vascular smooth muscle cells"

**Supplemental Figure 1**

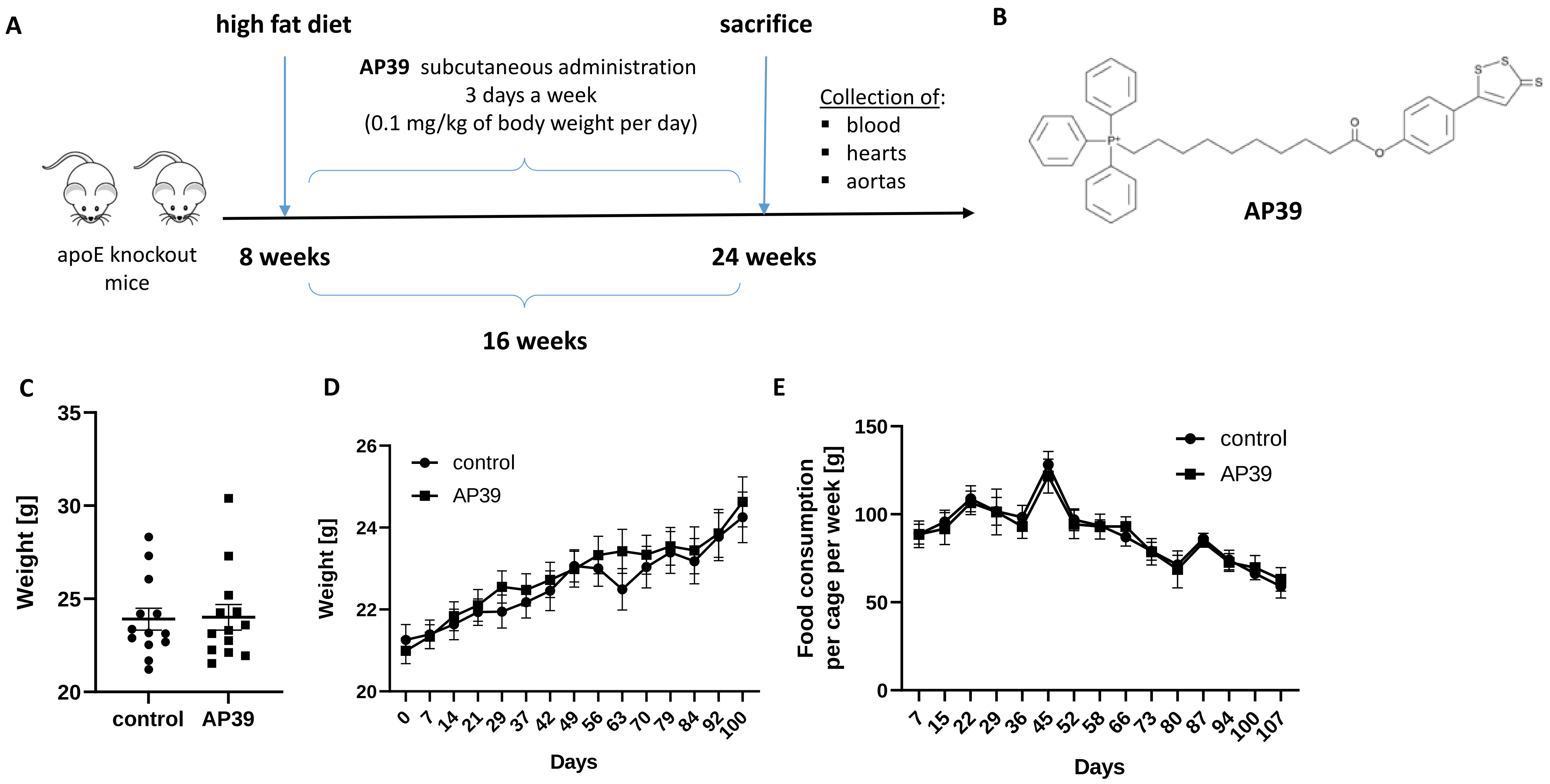

Supplemental Figure 2

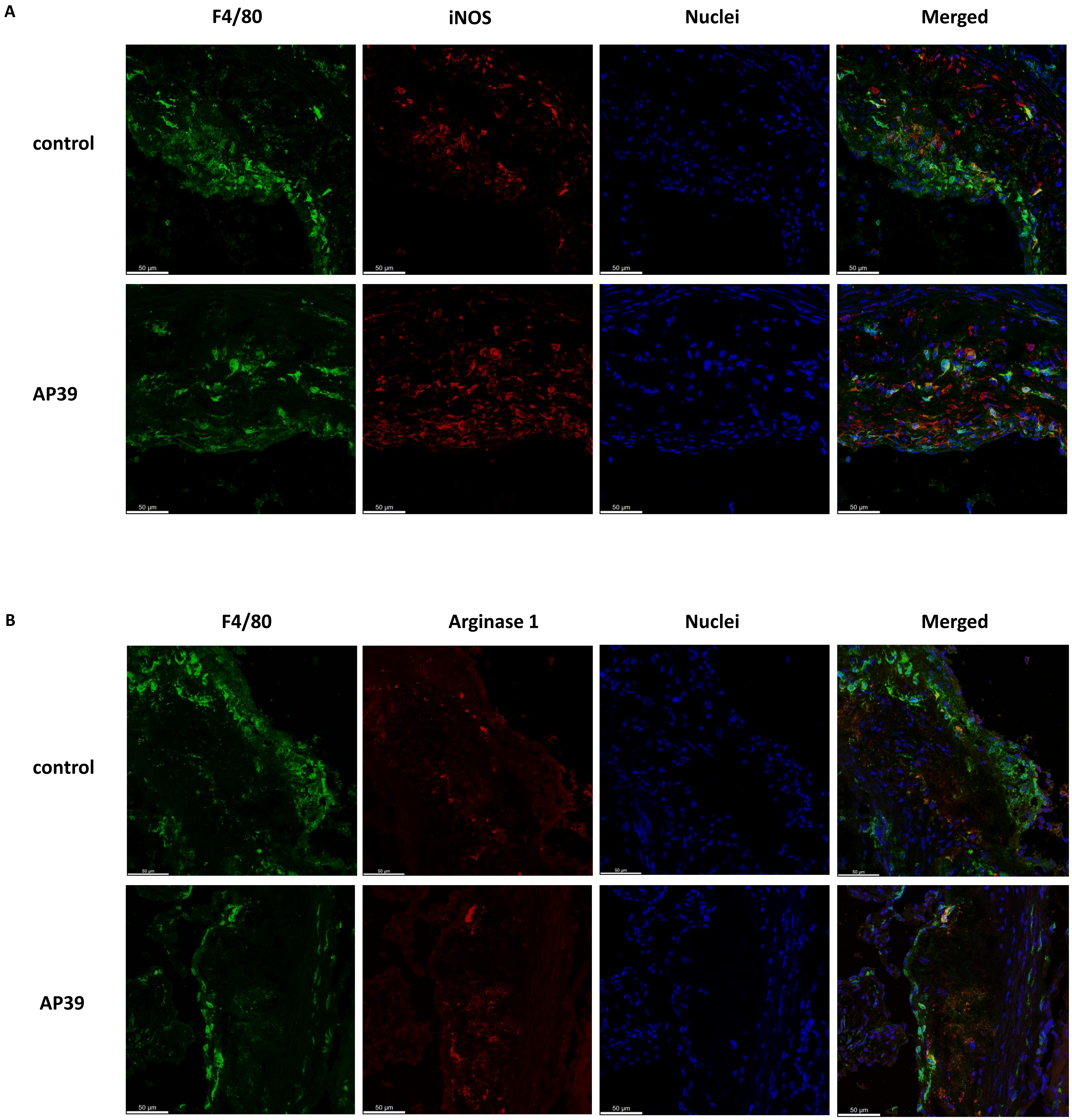

**Supplemental Figure 3**

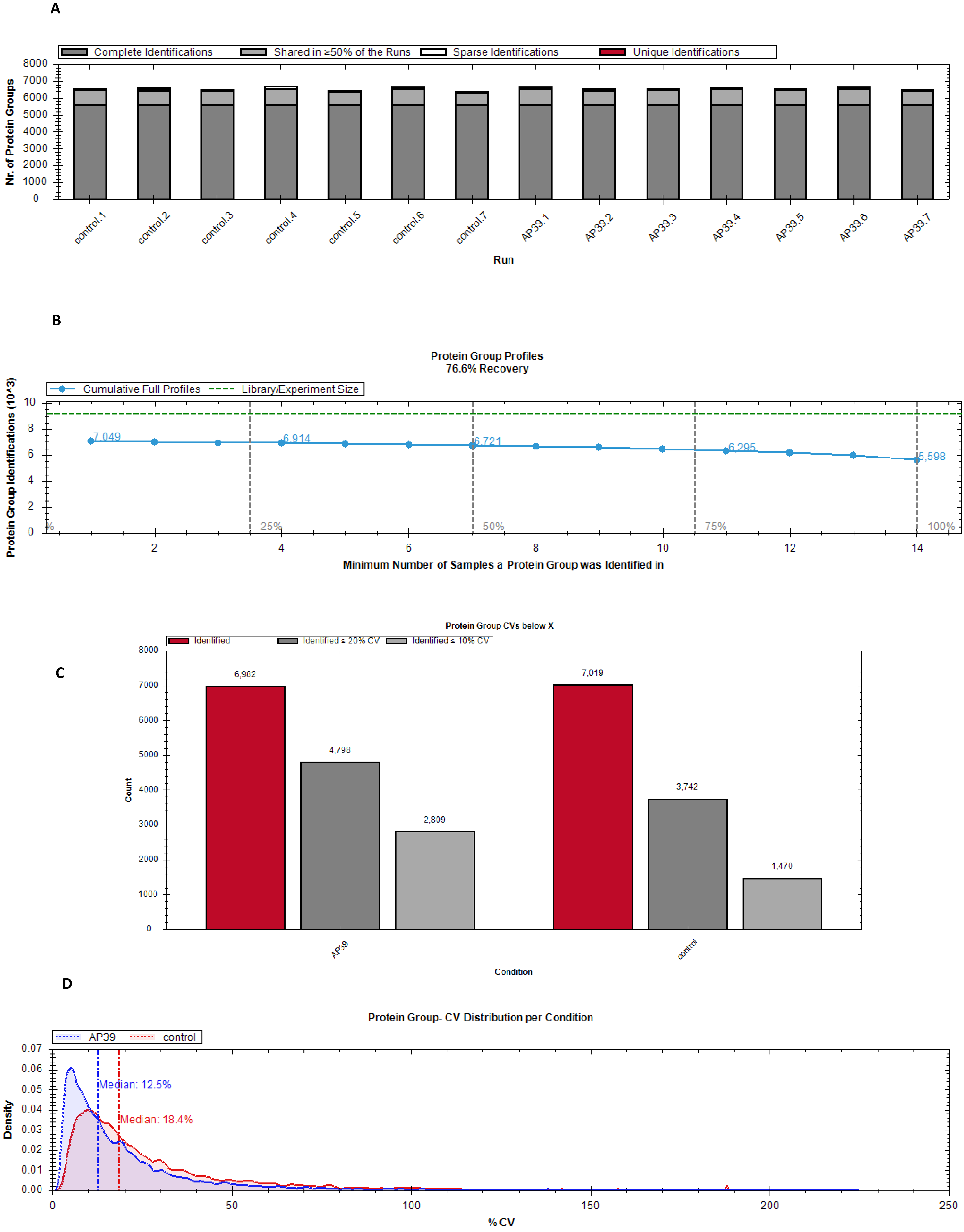

Supplemental Figure 4

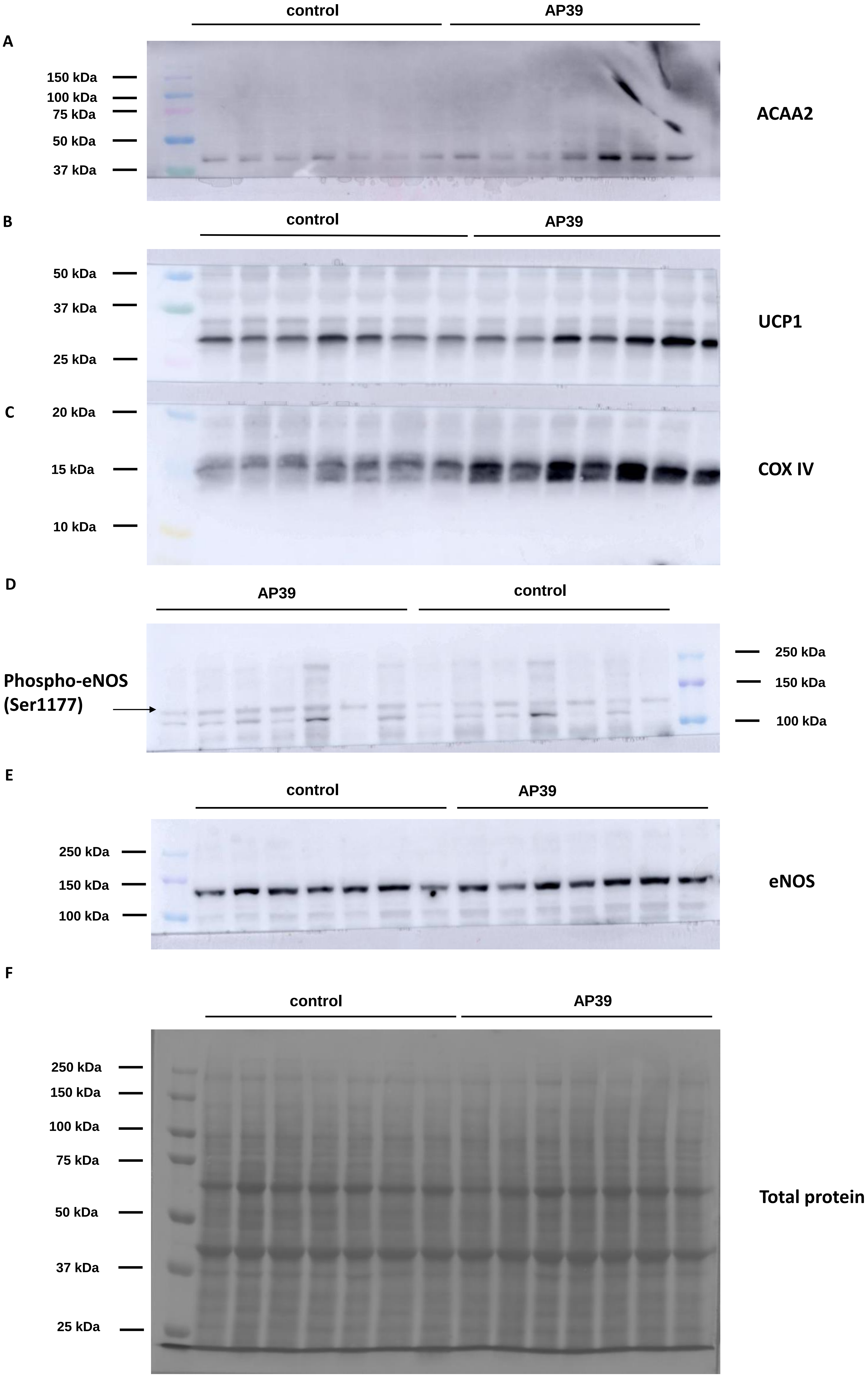

Supplemental Figure 5

A

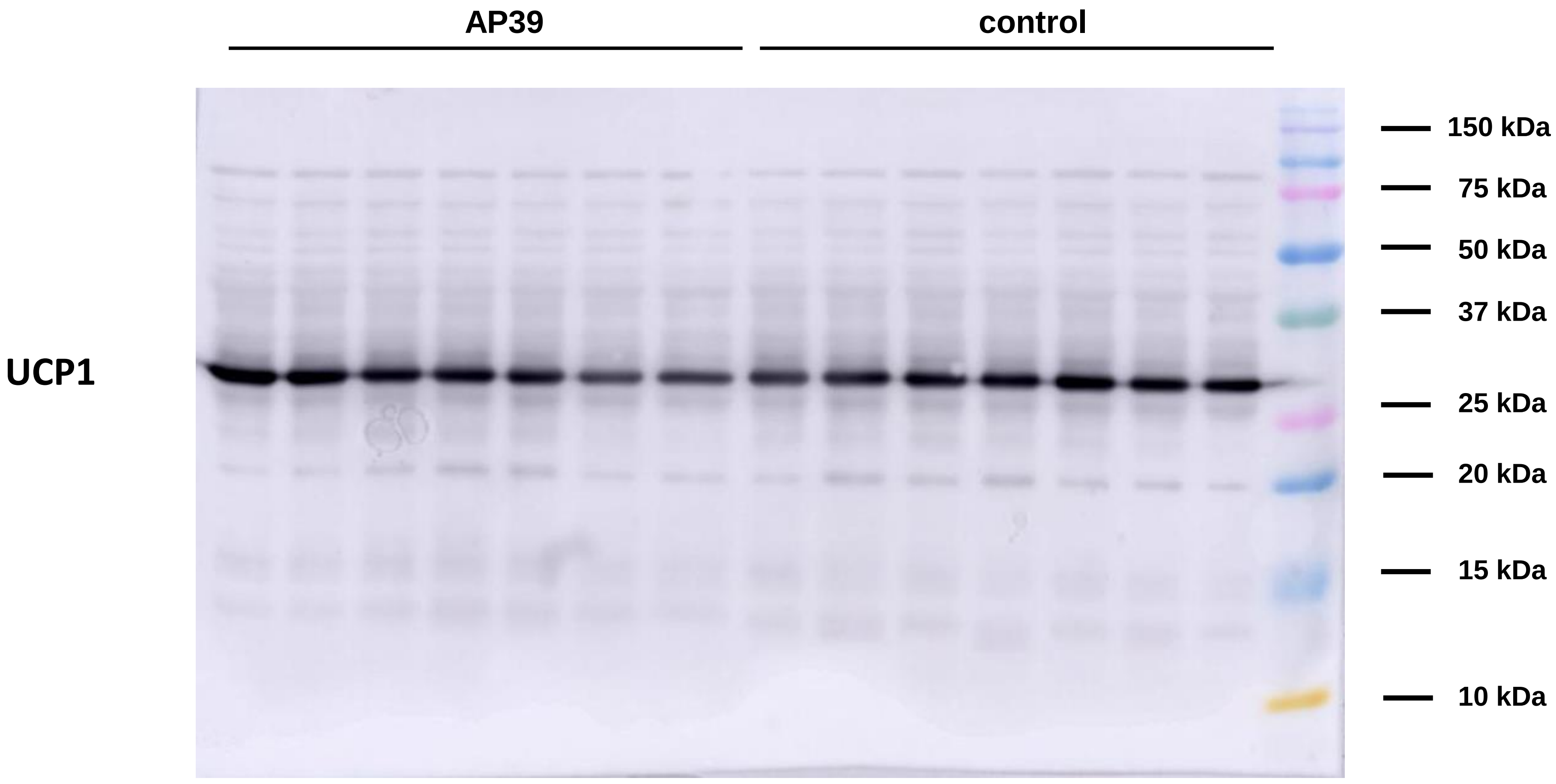


B

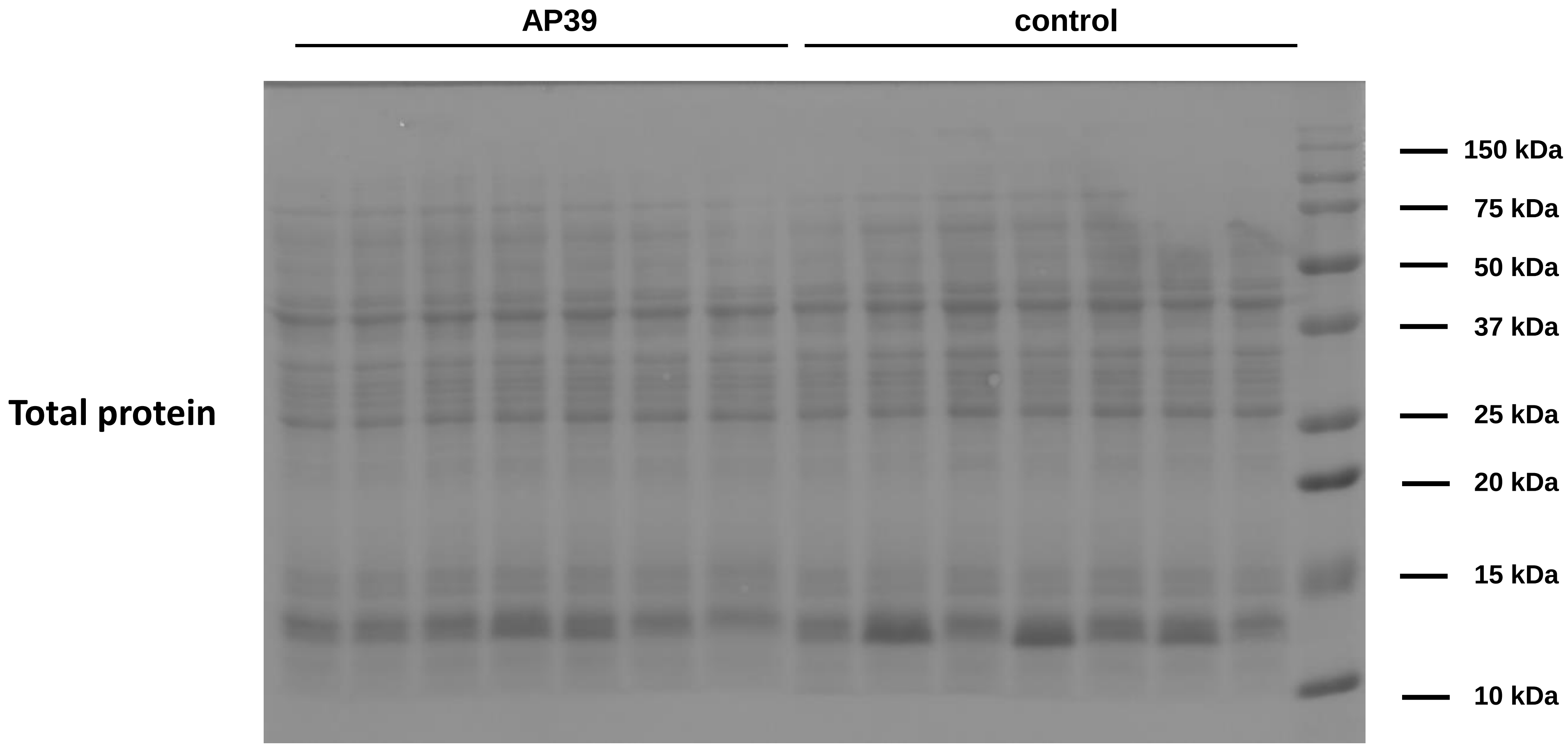

**Supplemental Figure 6**

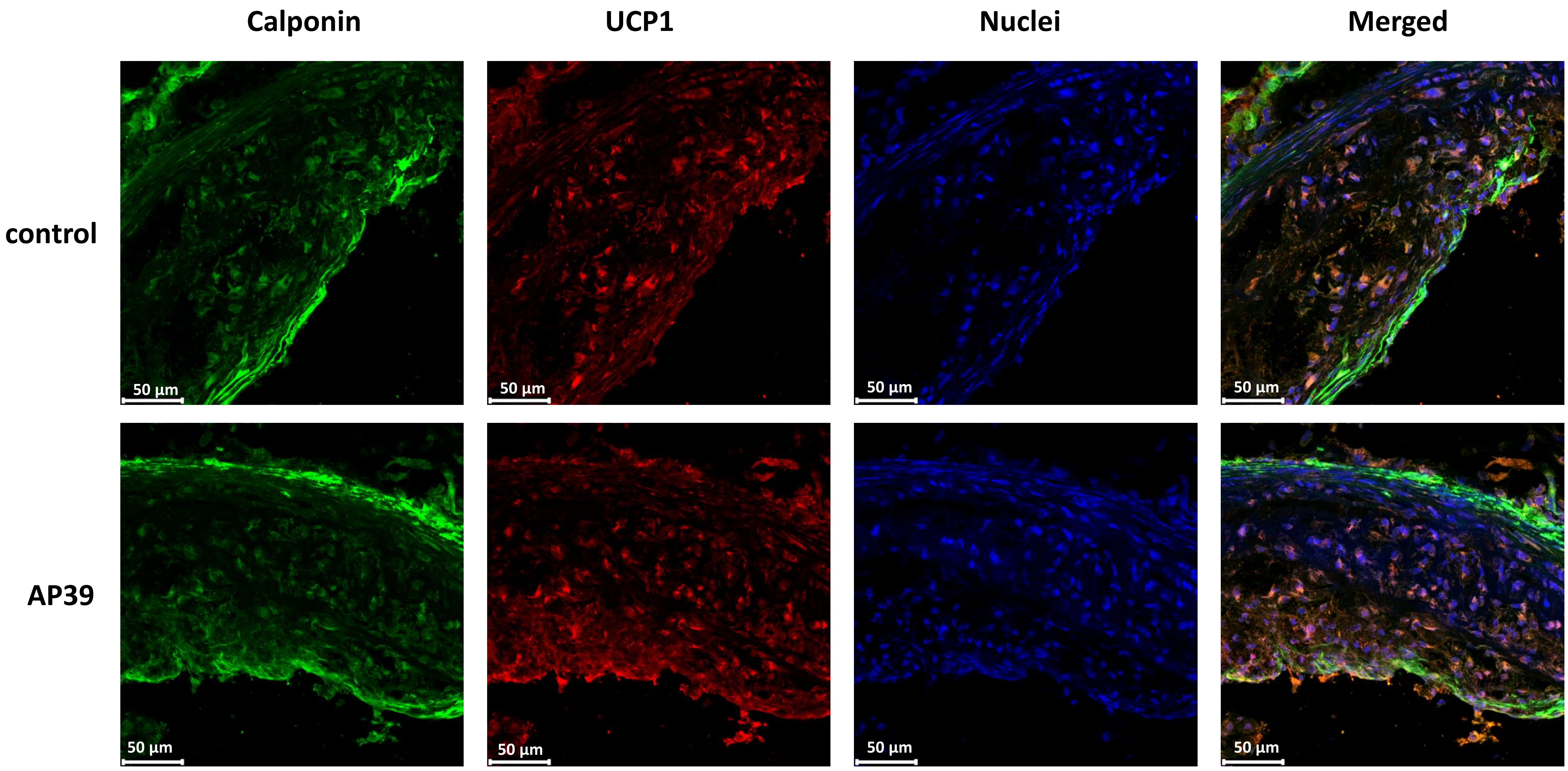

**<u>Supplemental Figure 1</u> The role of AP39 in atherogenesis - an experiment overview.** The experimental design of the study **(A)**. The chemical structure of AP39 **(B)**. Weight of control and AP39-treated mice at the end of experiment **(C)**. Weight of control and AP39-treated mice over 16 weeks of high-fat diet and drug administration **(D)**. Food consumption rate of control and AP39-treated mice over 16 weeks of high-fat diet and drug administration **(E)**. Mean ± SEM; n=13.

**<u>Supplemental Figure 2</u> Immunohistochemical stainings of M1 and M2 macrophages in atherosclerotic lesions.** Representative immunohistochemical staining of aortic roots showing F4/80 (green), 4’6-diamidino-2-phenylindole (DAPI) (blue) and nitric oxide synthase 2 (iNOS) (red) (M1 phenotype) (**A**) or arginase 1 (red) (M2 phenotype) (**B**) co-localization in control and AP39-treated mice.

**<u>Supplemental Figure 3</u> Quality control of MS runs of the aorta of control and AP39-treated mice.** Protein group identification details across all LC-MS runs **(A)**. Spectral library recovery **(B)**. Coefficient of variations (CVs) for protein groups across all biological conditions **(C)**. Distribution of protein group CV in biological conditions **(D)**.

**<u>Supplemental Figure 4</u> Western blot replicates as uncropped images. Related to Figure 4B, 5A and 7A.** Western blot replicates for 3-ketoacyl-CoA thiolase (ACAA2) **(A)**, mitochondrial brown fat uncoupling protein 1 (UCP 1) **(B)**, cytochrome C oxidase subunit IV (COX IV) **(C)**, phospho-endothelial nitric oxide synthase (p-eNOS) **(D)**, eNOS **(E)** in the aorta. Ponceau S staining used to normalize data for total protein level **(F)** (n=7).

**<u>Supplemental Figure 5</u> Western blot replicates as uncropped images. Related to Figure 5A.** Western blot replicates for mitochondrial brown fat uncoupling protein 1 (UCP 1) in PVAT **(A)**. Ponceau S staining used to normalize data for total protein level **(B)** (n=7).

**<u>Supplemental Figure 6</u> Immunohistochemical stainings of UCP1 in VSMCs in atherosclerotic lesions.** Representative immunohistochemical staining of aortic roots showing calponin (marker of VSMCs) (green), 4’6-diamidino-2-phenylindole (DAPI) (blue) and uncoupling protein 1 (UCP1) (red) co-localization in control and AP39-treated mice.
